# Contrastive learning of adverse events to provide effective and interpretable vector representations for machine-assisted pharmacovigilance

**DOI:** 10.1101/2025.06.04.657852

**Authors:** Olivér M Balogh, Mátyás Pétervári, Áron M Csernák, Eszter Puhl, András Horváth, Péter Ferdinandy, Bence Ágg

**Author notes:** correspondence: Bence Ágg, MD, PhD.

## Abstract

Post-marketing surveillance is crucial for drug safety, yet the tools of pharmacovigilance rely solely on text-based data that may limit the applicability of contemporary machine learning methodologies in the support of decision making. Here, we adapt contrastive learning algorithms to generate adverse event vector representations from spontaneous reports to serve as general machine-readable resources for pharmacovigilance applications. We present comprehensive analyses of the resulting representations through density-based clustering, semantic evaluation and comparison of multivariate dispersions, revealing patterns that reflect both functional and causal relations of the adverse events while also capturing drug-safety related information better than existing medical terminologies and encoder-only large language models (LLMs). Furthermore, we demonstrate the applicability of the representations as input features in our downstream model, outperforming the reporting odds ratio method commonly used by regulatory agencies (AUROC: 0.88 vs 0.75) and LLM-based representations (AUROC: 0.88 vs 0.83) on drug–event causality prediction benchmarks. As such, this is the first demonstration of an interpretable adverse event vector representation that can be utilized for training arbitrary models, enabling wider and more effective applications of machine learning in pharmacovigilance.

## 1. Introduction

Adverse events are any harmful medical occurrences experienced during the administration of a medicinal product. When suspicion is conveyed about a possible causal relationship between an adverse event and a product, the terms adverse effect, adverse reaction or adverse drug reaction (ADR) are used instead^1^.

The role of pharmacovigilance is to minimize the risk-to-benefit ratio of drugs via monitoring their adverse reactions throughout both the development and post-marketing phases. Several adverse reactions appear only during the latter, recorded in spontaneous adverse event reports (AERs). AERs are collected for analysis by various pharmacovigilance entities, such as the EudraVigilance system of the European Medicines Agency (EMA)^2^, the Adverse Event Reporting System of the United States Food and Drug Administration (FAERS)^3^, and the VigiBase database of the World Health Organization (WHO)^4^. AERs suggest only the potential for causality between drugs and adverse events (or equivalently in this case, adverse reactions), and thus they require careful and time-consuming causality assessment by experts before the appropriate regulatory action can be issued. Given the sheer volume of AERs (e.g. more than 2 million received by FAERS in 2023^5^), there is an increasing need for efficient automation methods to either improve the identification of new safety concerns from the reports that are likely caused by drugs (a process called signal detection) or to accelerate the causality assessment of already suspected drug–event pairs. Drug safety professionals have expressed their keenness to engage with artificial intelligence systems in order to shift their volume-based tasks to a value-based work^6–10^, and so, our aim here is to provide a general, interpretable resource for signal detection and causality assessment in machine-assisted pharmacovigilance, both for applications and future research.

A core limitation of handling AERs is their inherent text-based structure, lacking any informative descriptor of the suspect medicinal products and the accompanying adverse events beyond their names. For drug substances, several chemical and biological descriptors are available from external databases, such as DrugBank^11^ or PubChem^12^, that can provide numeric features for machine learning. There is, however, no similarly rich, machine-readable resource for adverse events. Representation learning could serve as a substitute by producing information-preserving embedding vectors, yet, to the best of our knowledge, no such data specifically made for general pharmacovigilance applications are readily available.

Representation learning of medical concepts is, however, not a new field of study. In terms of embedding style, we can identify: (i) methods with traditional word2vec-like approaches^13–21^, (ii) methods where knowledge graphs or term hierarchies are also constructed, resembling graph embeddings^21–25^, (iii) and methods that are based on pretrained encoder-only large language models (LLMs), such as BERT^25–32^. Regarding the application of these models in pharmacovigilance, one of the most frequent obstacles is that they embed on the level of various Unified Medical Language System subsets^15,16,18,19,21,25^ with only a partial mapping to the preferred terms (PTs) of the Medical Dictionary for Regulatory Activities (MedDRA^®^), which is the most commonly used terminology standard in pharmacovigilance^33^. For static embeddings (i.e. word2vec-like and graph-based ones), this obstacle introduces either a complicated task of needing to translate between the terminologies, or significantly limits the number of embedding vectors readily available for pharmacovigilance pipelines. For contextual embeddings produced by encoder-only LLMs, this problem is bypassed through the subword- or character-level tokenization of out-of-vocabulary words and the use of attention. However, LLMs are rarely utilized to provide individual word (or term) embeddings in a static fashion, as the methods to do so (e.g. context aggregation, decontextualization) have their own challenges and limitations, especially in terms of interpretability^25,34,35^. The second recurring aspect hindering the reuse of medical concept embeddings is that they are often learnt through a very specific, non-generalizable task and are part of a larger end-to-end prediction model, with no description of their applicability in other downstream models. For example, encoder-only LLMs fine-tuned for language tasks, such as document classification, natural language inference and named entity recognition, all belong to this category^26–32^, as well as some of the word2vec-like models^17^ and graph-based embeddings^22,24^. The third notable limitation of reusing medical embeddings in pharmacovigilance is that most of them utilize natural text as their input, ranging from PubMed abstracts and full text articles^15,16,19,21,27–31^ to clinical notes and electronic health records^15,16,26,29,30^ to even insurance claims and social media posts^15,16,18,28^, making AERs an underutilized resource for representation learning, despite their relevance in pharmacovigilance. Lastly, availability is also an issue as some embeddings have simply not been made public^14,20^.

Motivated by these limitations of previous studies, we created our own representation learning framework that is based on a data processing module (Fig. 1a), handling raw FAERS reports, and an embedding module (Fig. 1b) responsible for learning the vector representation, i.e. embedding, of adverse events (from here on simply referred to as reactions) on the level of MedDRA^®^ PTs. In order to investigate a more diverse set of embedding methods, we developed two models: one based on normalized temperature-scaled cross entropy^36^ (here referred to as NTX), and one based on the negative sampling skip-gram version of word2vec^37^ (here referred to as NSG). In both cases, the embeddings are learnt via noise-contrastive estimation^38^ for which the reactions are first paired up with their context terms as observed in the AERs. We defined three different context-sampling approaches based on what is considered as context for a given “target” reaction: only drugs it was reported with (reaction2drug), only the other reactions it was reported with (reaction2reaction), or both (reaction2all), resulting in six different embedding sets overall. Comparing this framework to learning embeddings from natural text, AERs are treated analogous to the sentences of a corpus, and our context-sampling approaches function like a skip-gram sampling method with “dynamic” context window that operates within only the current sentence. However, in our framework, the distribution of context terms is influenced by the drug-safety related behavior, i.e. reporting patterns, of the given reaction, rather than its occurrences in written language. Consequently, the resulting embeddings encode the pharmacovigilance profile of reactions, instead of their linguistic characteristics, which makes these embeddings not only effective descriptors for downstream applications but also semantically interpretable through various analysis methods.

**Figure 1.**
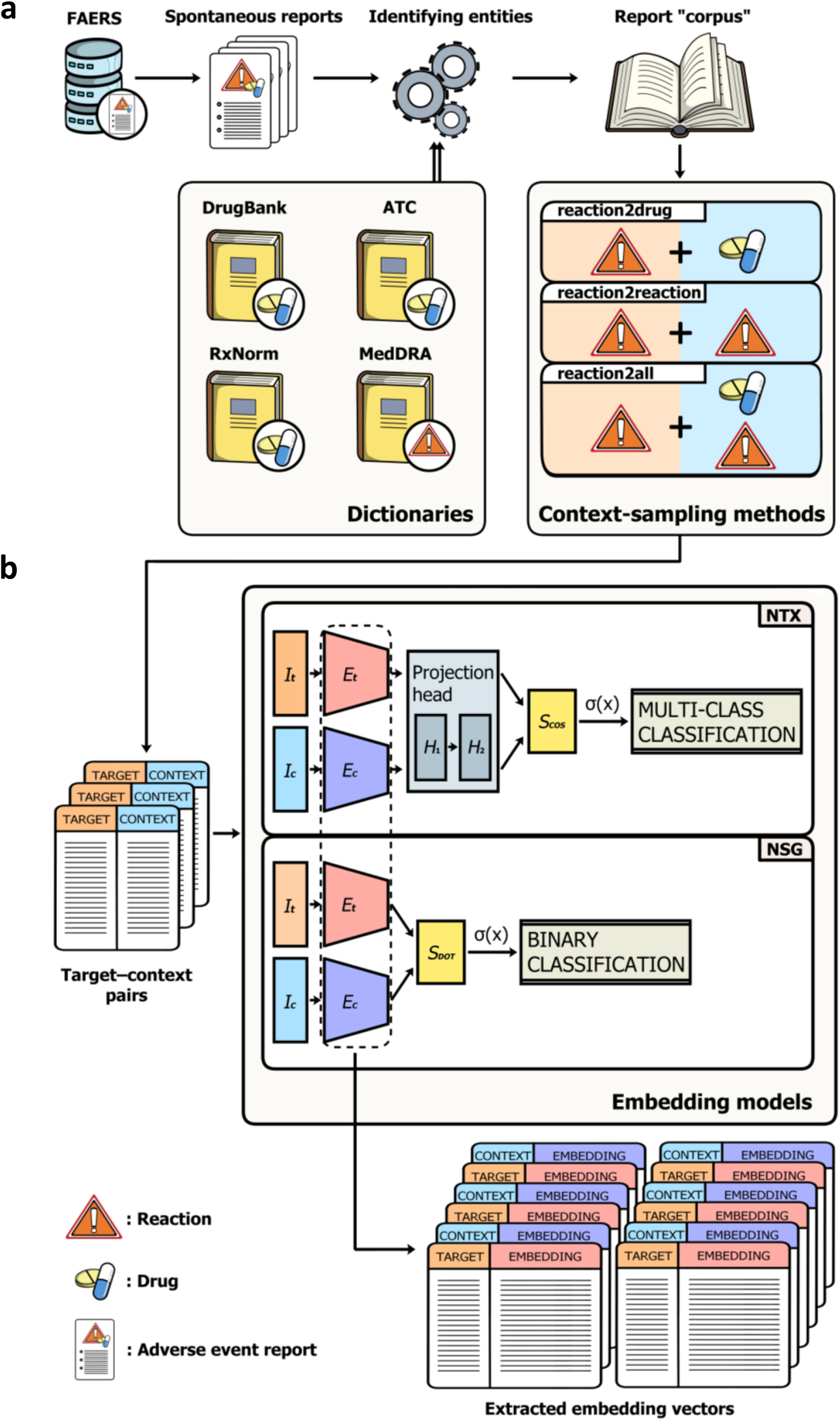
Schematic visualization of the main framework. **(a)** Data processing: we accessed spontaneous adverse event reports from the FAERS database, and used a combination of various naming dictionaries to identify the suspect drugs and reactions, mapping them into canonic DrugBank substance names and MedDRA^®^ preferred terms, respectively. This step also acts as a quality filter. We performed three different context-sampling methods on the resulting report “corpus”, based on what is considered as context for a given reaction: the drugs it was reported with (reaction2drug), the other reactions it was reported with (reaction2reaction), or both (reaction2all). **(b)** Embedding: the three types of target–context pairs were utilized as input data for our embedding models, which learn the vector representation, i.e. embedding, of the reactions through two different contrastive learning algorithms. The NTX model is based on the normalized temperature-scaled cross entropy loss, and performs multiclass classification on batches of target–context pairs, where each target is expected to be classified as belonging only to its corresponding context and vice-versa. The NSG model uses negative sampling, and performs binary classification on the target–context pairs (positive class) that are complemented with randomly sampled negative pairs (negative class). In both cases, the embedding models are discarded after training, and only their embedding matrices are kept for further analysis.

## 2. Results

To characterize the interpretability and effectiveness of the reaction embeddings, we conducted three different analyses, consisting of visualizations (Fig. 2a), semantic (Fig. 2b) and statistical (Fig. 2c) evaluations and a use case demonstrating real life applicability for drug–event causality prediction (Fig. 2d). Although separate embeddings are learnt for target and context terms due to the structure of the models (Fig. 1a), we only focus on the target ones, which is common practice due to the learning processes (i.e. term distribution adjustments and training data sampling) being primarily designed around optimizing the target embedding vectors.

**Figure 2.**
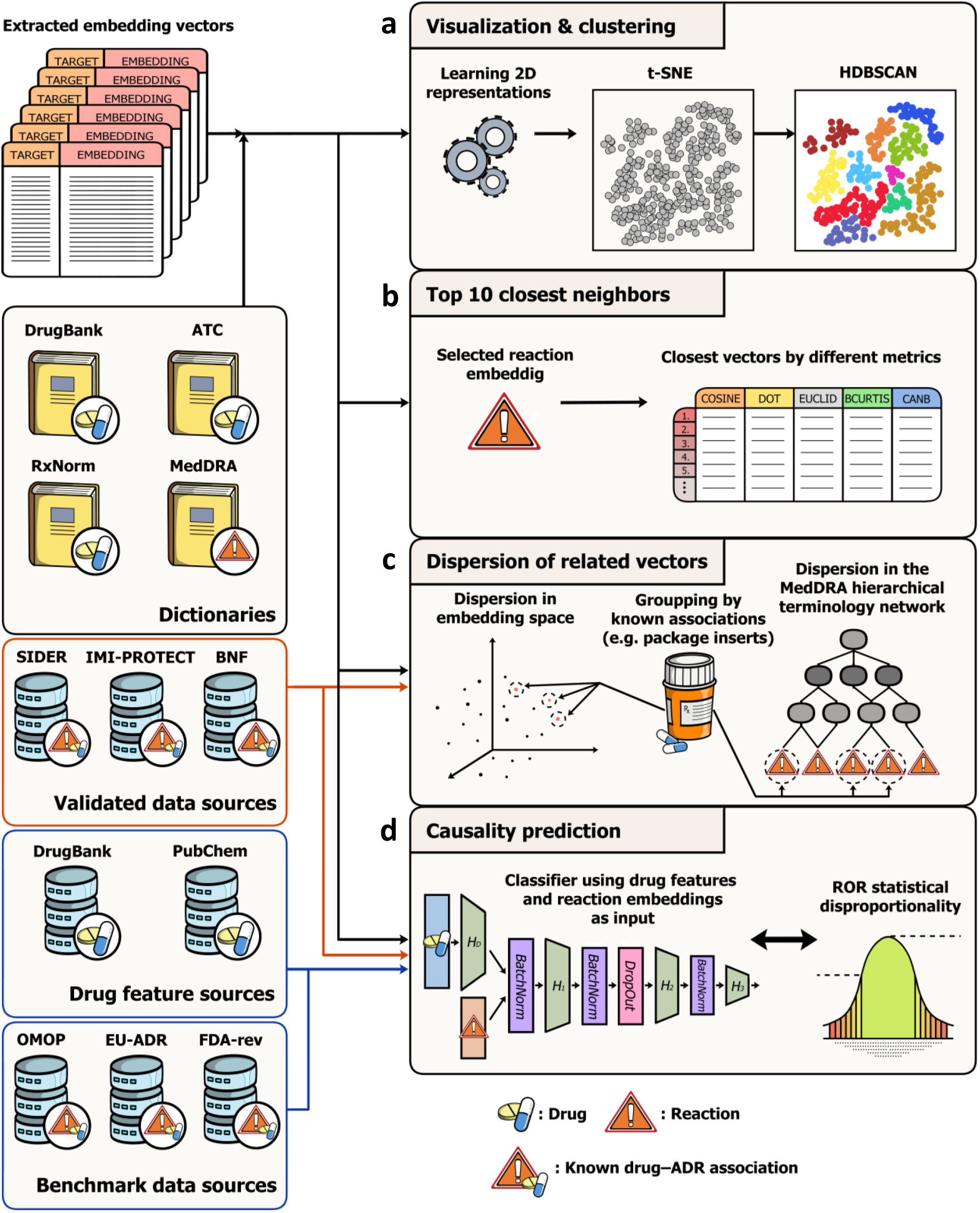
Summary of our analyses methods and the data sources used to characterize the reaction embeddings. **(a)** We applied t-SNE for dimensionality reduction and HDBSCAN clustering to support the visualization by discovering potentially meaningful clusters of reactions. **(b)** By calculating various distance and similarity measures in the embedding space, we evaluated the semantic similarities of top 10 closest neighbors of a selected reaction. **(c)** Using validated drug– ADR (adverse drug reaction) data, we analyzed the multivariate dispersion of embedding vectors for reactions that belong to the same drug. Dispersions of validated groups were statistically tested against randomly selected groups, and also compared to the dispersions measured in the MedDRA^®^ hierarchical terminology network. **(d)** We constructed a deep neural network classifier to utilize the embedding vectors as input features, along with drug features from DrugBank and PubChem, for drug–event causality prediction. The model was trained on the validated drug–ADR sources with various negative sampling approaches (Section 4.9), and its performance was evaluated on established benchmark sets, comparing it with using embeddings based on large language models and the reporting odds ratio (ROR) statistical disproportionality analysis method.

### 2.1. Visualization and clustering

Visualizing the embeddings enables us to gain a general intuition about the underlying data structures. With the inherent limitations kept in mind, we applied *t*-distributed stochastic neighbor embedding (*t*-SNE)^39^ to reduce the original representations into 2D. For a deeper analysis, we selected the NSG-reaction2drug embedding (see Supplementary Fig. 1-5 for the others). The reaction2drug-based embeddings have a perspicuous explanation, i.e. vectors will be close to each other for reactions that were reported with the same or similar drugs. Out of the two embedding models, NSG produced the visually more interpretable reaction2drug representation, for which a logarithmic frequency-based coloring (Fig. 3a) revealed a further, yet evident property: during training, the NSG model encountered the often reported reactions much more frequently, even with subsampling applied, thus learning embedding vectors for them that are a lot more refined and pulled farther away from the randomly initialized values. We can observe the most frequently reported reactions congregating in the middle of the plot and on the centre-facing edges of most of the visually well-separated clusters. These are general terms, such as *Drug Ineffective, Pain, Off Label Use, Fatigue, Nausea* and *Death*. At the top-left side of Fig. 3a, we can also observe a large group with low frequency, lacking a clear set of frequent reactions to gravitate toward. This group holds embedding vectors that had such a low occurrence during training that they barely moved from the noise initialization. Here, we have a diverse repertoire of niche reactions, such as *Vaccine Breakthrough Infection, Benign Rolandic Epilepsy, Cholesteatoma Removal, Mechanic’s Hand, Perthes Disease* and *Coccidioides Encephalitis*.

**Figure 3.**
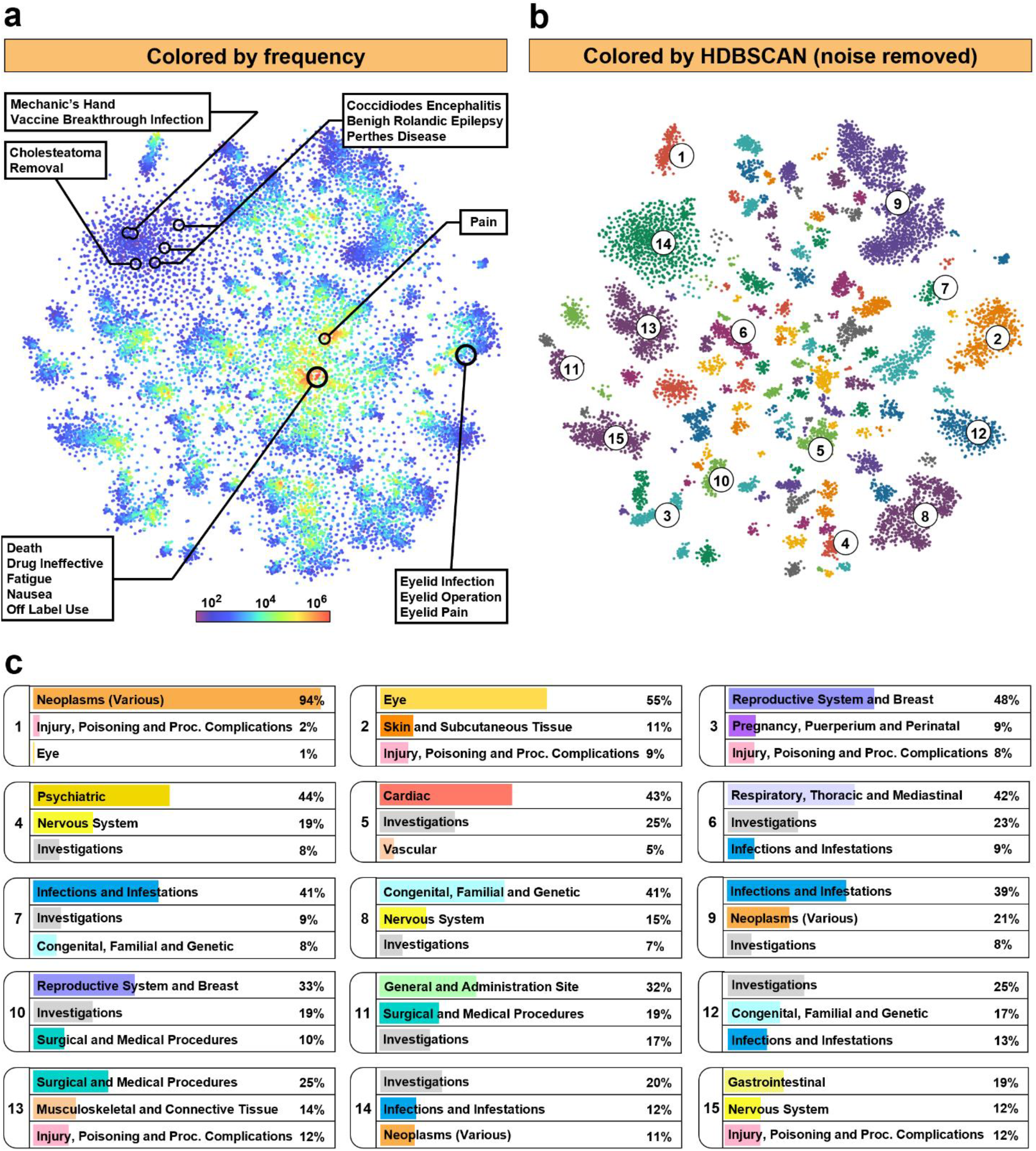
t-SNE visualization of the embedding space generated by the NSG model with reaction2drug context-sampling. **(a)** Frequency-based coloring of the t-SNE plot, using occurrence counts from the processed adverse event reports with a logarithmic scale for smoother color transitions. Example reactions discussed in Section 2.1 have their positions indicated. **(b)** HDBSCAN-based coloring of the t-SNE plot, with noise removed for a better visual separation. **(c)** Ratio of the top three MedDRA^®^ System Organ Classes (SOCs) in each selected cluster. Selected clusters are numbered in descending order of their highest SOC ratio. SOC names are shortened to fit the figure by leaving out the words “disorders” and “conditions”, as well as the list of neoplasm types.

Supporting our visualization are the clusters deemed most prominent by Hierarchical Density-Based Spatial Clustering of Applications with Noise (HDBSCAN) (Fig. 3b), which we chose for its robust performance on variable densities and ability to prune off noisy elements^40^. Small-scale density differences are not preserved by *t*-SNE, but with the elimination of noise, HDBSCAN still proved successful in highlighting well-separated clusters for further analysis. For example, we can examine the composition of highest level MedDRA^®^ categories, called System Organ Classes (SOCs), in the resulting clusters to get an approximate idea how closely the embedding space resembles them (Fig. 3c). Given that there are 27 distinct SOC terms, we selected only the top three with the highest ratios in each cluster, and chose 15 clusters out of 127 in total, focusing on the largest and most representative ones. The first cluster is almost homogeneous, filled with reactions related to tumors (neoplasms), while the other clusters are more heterogeneous, with a pronounced plurality class. This is unsurprising, as reactions related to each other on the basis of prescribed therapies (and evidently, anatomical considerations) are often delegated into different SOCs, which is a phenomenon most prevalently caused by the umbrella SOCs, like those relating to investigations, injuries, infections and procedures (Fig. 3c). For example, *Eyelid Pain* is categorized by MedDRA^®^ as being primarily an eye disorder, while *Eyelid Infection* is an infection and *Eyelid Operation* is a surgical procedure. Yet, the strong relation of these terms is reflected by the embedding space, with all three appearing close to each other in the second cluster, despite the embedding model lacking word and subword information (e.g. eyelid) due to the term-level “tokenization”. This shows that the embedding space has a clear, interpretable structure that deviates from MedDRA^®^ categories, but various anatomically and functionally consistent groups still emerge on the macro-scale. The local patterns, on the other hand, retain high similarities with MedDRA^®^, as exemplified in Section 2.2., while the deviations in global structure are demonstrated to capture drug-safety related information better than MedDRA^®^ in Section 2.3.

### 2.2. Top 10 closest neighbors

Here we present a brief semantic analysis through the example of a MedDRA^®^ PT, *Myocardial Ischaemia*, and its 10 closest neighbors across the representations to provide a closer insight into the local patterns of the embedding spaces without resorting to dimensionality reduction (Fig. 4, Supplementary Table 1-7). Ischaemic heart disease (MedDRA^®^ PT: *Coronary Artery Disease*), is a major health concern, being ranked as the most prevalent cardiovascular illness and a leading cause of death worldwide^41^. It involves the buildup of partial or complete obstruction in the coronary arteries (*Arteriosclerosis Coronary Artery*), resulting in the development of *Myocardial Ischaemia*, a condition that is characterized by a mismatch of oxygen demand and blood supply (*Ischaemia*) in the cardiac muscle. This induces metabolic deficiencies at the cellular level, eliciting progressive tissue damage and, consequently, death of myocardial tissue (*Acute Myocardial Infarction*), heart rhythm abnormalities (*Arrhythmia*) and impaired pump function (*Cardiac Failure*). We selected *Myocardial Ischaemia* as the focus of this evaluation because of its relevance to global research^42^, rich literature^43^ and connection to our previous studies^44,45^. By our provided code, the list of top closest neighbors can be easily accessed for any other reactions as well (see Code availability).

**Figure 4.**
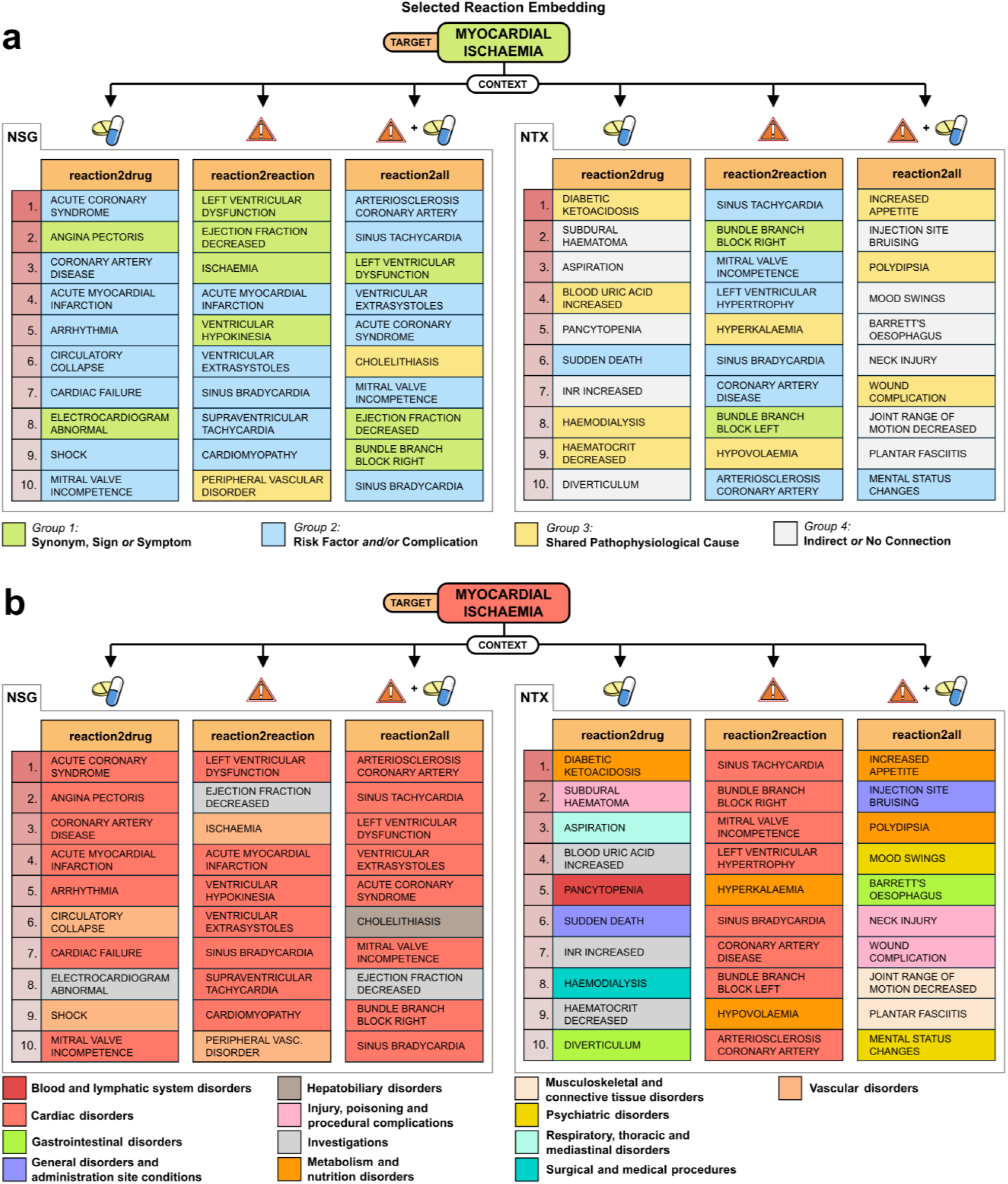
Lists of the top 10 closest neighbors of the MedDRA^®^ preferred term Myocardial Ischaemia across the six reaction embeddings, measured by cosine similarity in the original vector space, colored **(a)** by semantic evaluation based on expert knowledge and literature, and **(b)** by primary MedDRA^®^ System Organ Class (SOC).

Pathophysiological connections and clinical relevance of the neighboring reactions of *Myocardial Ischaemia* were assessed via international medical guidelines (Supplementary Table 8), categorizing them into one of four distinct groups (Fig. 4a) with MedDRA^®^ SOC categorization as reference (Fig. 4b). Group 1 consists of synonyms, such as the aforementioned *Ischaemia*, and the various signs/symptoms considered during the diagnosis of *Myocardial Ischaemia* itself, such as *Angina Pectoris* (chest pain) and *Electrocardiogram Abnormal*. Group 2 contains the most neighbors, corresponding to various pathophysiological risk factors, like *Arteriosclerosis Coronary Artery* or *Sinus Tachycardia/Bradycardia* (heart beats faster/slower than normal), and possible complications of *Myocardial Ischaemia*, such as *Arrhythmia* or *Cardiac Failure*. For Group 3, we selected reactions that share their pathophysiological cause with *Myocardial Ischaemia*, and have identifiable common risk factors other than old age. For example, both *Peripheral Artery Disease* and *Myocardial Ischaemia* can be caused by arteriosclerosis, with diabetes mellitus, hypertension and dyslipidaemia (abnormal levels of lipids in the blood) being well-known risk factors. Finally, Group 4 consists of reactions with semantic connections that are too indirect to be relevant.

We can observe several different tendencies between the embeddings of the two models. NSG is consistent in emphasizing closely connected concepts across the three context-sampling methods, with distinguishable preferences among them: reaction2drug prioritizes the most well-known complications and symptoms, reaction2reaction leans toward terms describing impaired pump function and heart rhythm problems, and reaction2all results in a balanced set of the two. In contrast, NTX embeddings resemble such patterns only with reaction2reaction, while the other two sampling methods lead to several terms that share pathophysiological causes with *Myocardial Ischaemia*, like *Polydipsia* (excessive thirst) or *Diabetic Ketoacidosis* (acidification of blood due to lack of insulin), which are typical symptoms of untreated diabetes mellitus. Furthermore, many indirect entries appear as well, suggesting patient-specific connections that are less based on pathophysiology and more on circumstantial factors. For example, *Subdural Haematoma* (bleeding in the skull) might occur from falling or collapsing caused by haemodynamic instability, and *Aspiration* is common in patients that require intensive care due to an acute exacerbation of *Coronary Artery Disease*. Certain unusual entries, such as *Diverticulum* (intestinal wall abnormality) or *Joint Range of Motion Decreased* might be due to their frequent co-occurrences with *Myocardial Ischaemia* in the elderly.

Overall, this analysis demonstrates how pathophysiological and clinical properties of various diseases (reactions) can be captured purely from pharmacovigilance data by our proposed representation learning framework, encoding them in local neighborhoods. Furthermore, this evaluation also characterizes the potential strengths and weaknesses of the different models with the different types of context-sampling approaches, indicative of their behavior in downstream applications, as shown in Section 2.4.

### 2.3. Dispersion of related reactions

So far, we observed that the embedding vectors tend to cluster differently than MedDRA^®^ SOCs, yet their immediate neighbors display strong semantic relations. For pharmacovigilance purposes, we would expect the representations to also preserve drug-safety related information, which we measure by using curated datasets of drug–ADR associations^46–48^, considering them as pairs in which the causal relationship is validated (from here on referred to as validated data). A fair assumption would be that reactions caused by the same drug (a “validated group” of reactions) share more similarities than unrelated reactions (a “random group”), thus should be more closely packed together in the embedding space (Fig. 1c). Calculating pairwise interpoint distances of reactions inside the validated groups and equal-sized random groups allows us to conduct a permutation test for homogeneity of multivariate dispersions^49,50^. In total, we retrieved the ADRs of 1619 drugs, out of which 1612 had more than one ADRs, enabling dispersion calculations. The trivial result, i.e. that the ratio of significantly different validated groups is above 0.95 with almost all distance measures for all embeddings (Supplementary Table 9), motivated us to switch to a finer measure of dispersion differences by employing average pairwise interpoint distances directly^51^. Here, we refer to this measure as normalized gain, or “gain norm” for short (Fig. 5a). We expect the gain norm to be large for embedding spaces that well preserve drug-safety related information (i.e. a large separation of validated groups from background noise), therefore we also utilized it to tune the hyperparameters of our embedding models, most importantly the number of epochs (Fig. 5b, Supplementary Fig. 6-10).

**Figure 5.**
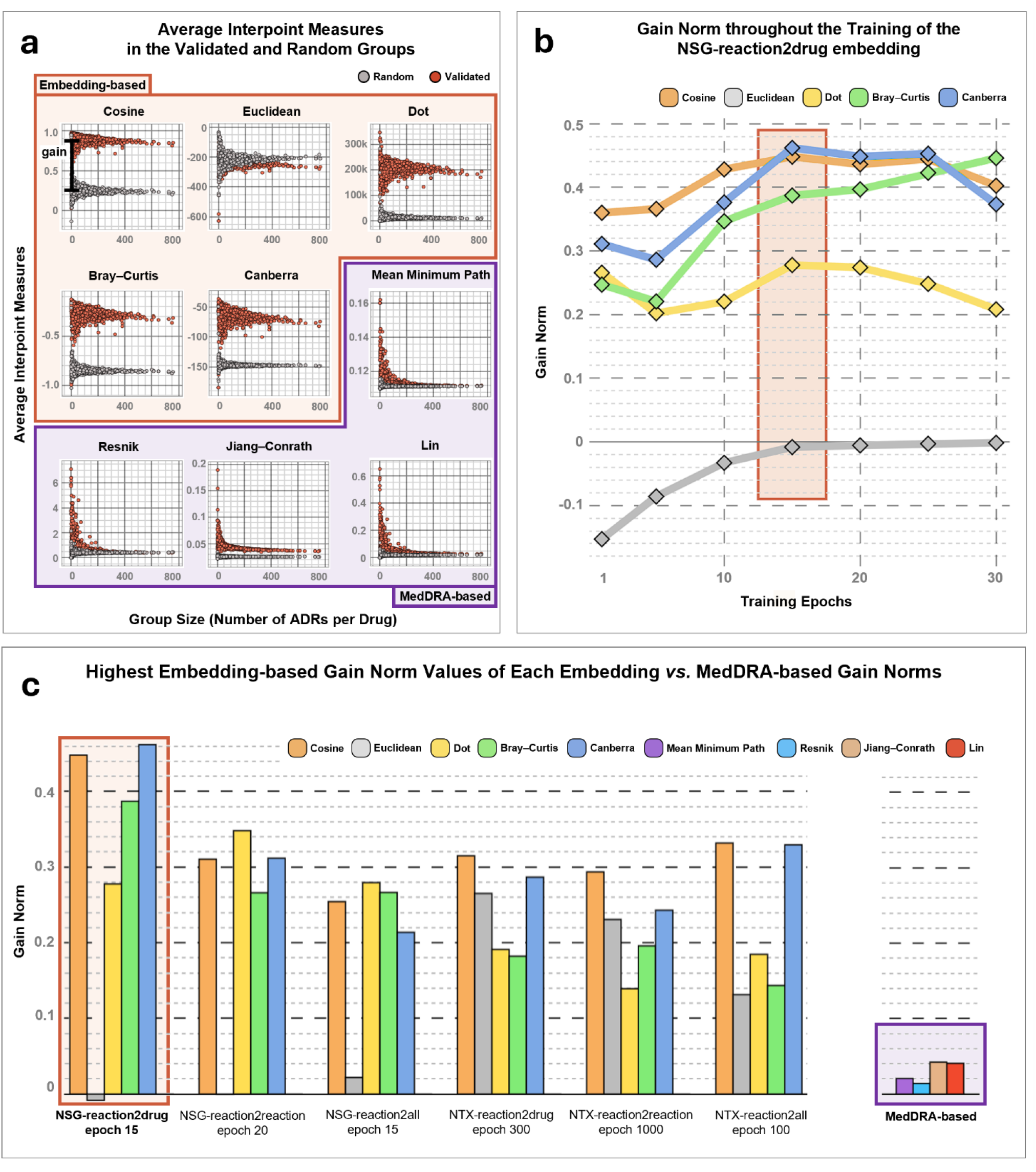
Representation of how the “gain norm” was calculated and used. **(a)** The concept of (unnormalized) gain: for each drug, we collected their validated reactions from known drug–ADR (adverse drug reaction) association data (validated group), and randomly sampled an equal number of other reactions (random group). The within-group average interpoint measures are plotted for both group types, as a function of group size, where the scale of the Euclidean, Bray– Curtis and Canberra distances are negated so that a higher value on the y-axis means more closely related groups on all plots. Gain is calculated as the average difference between the validated and the random groups. **(b)** Changes in the gain norm (normalized gain) as seen throughout the training of the NSG-reaction2drug embedding. We utilized the gain norm as a heuristic method for selecting the best epoch for each embedding. Highlighted here is the epoch selected for NSG-reaction2drug, from which data is visualized in panel (a) as well. **(c)** Summary of the highest scoring embedding results and their embedding-based gain norm values. Highlighted are the results belonging to the NSG-reaction2drug embedding, used as an example in panel (a) and (b). MedDRA-based gain norms were calculated by using hierarchical similarity measures on the MedDRA^®^ terminology network, which is independent of the embeddings, and so the results are summarized separately.

**Figure 6.**
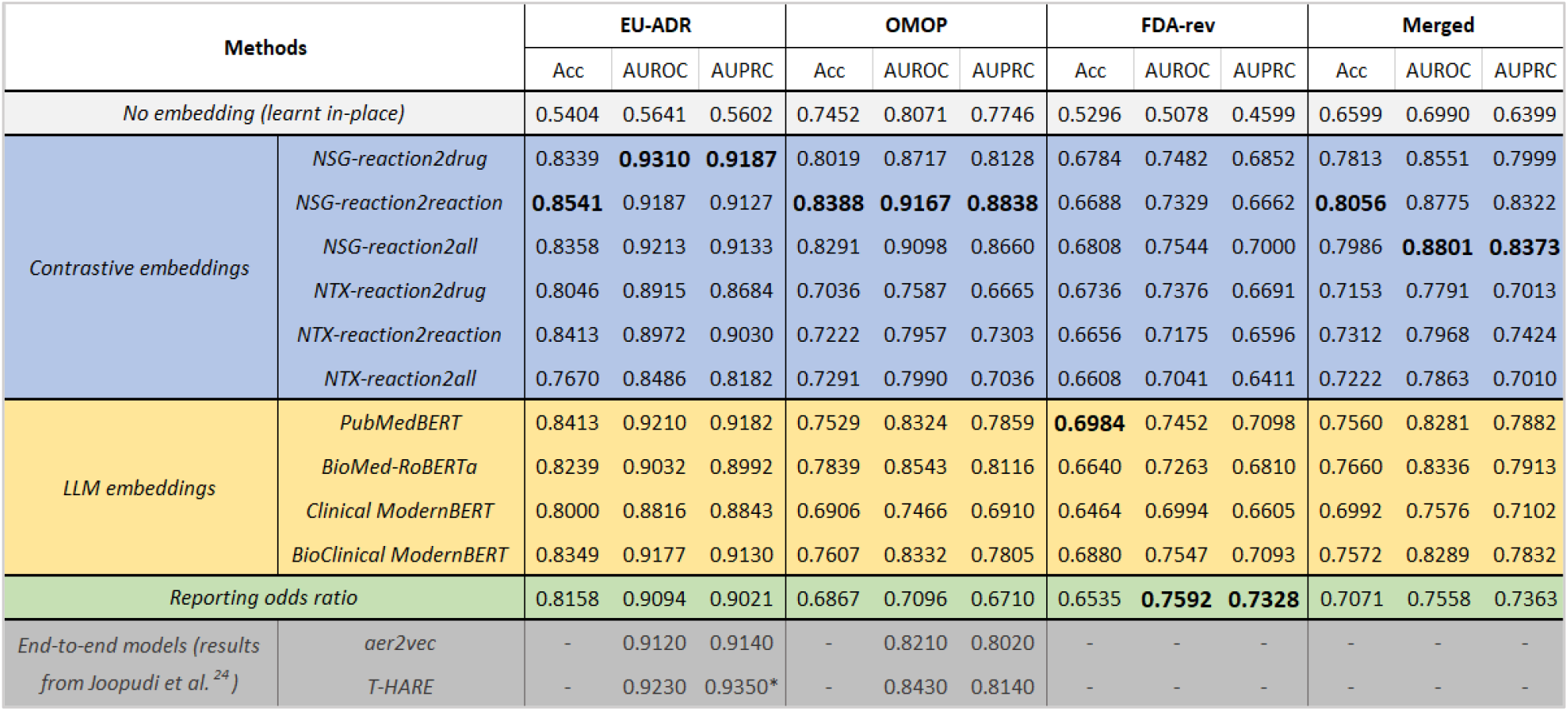
Drug–event causality prediction results. Performance of the classifier was tested by training it with no embeddings (learning them in-place), with using the presented six contrastive embeddings, and with using the four large language model (LLM) generated embeddings. As an additional baseline, the performance of the reporting odds ratio signal detection method was also calculated, and for further comparison, the results of two more end-to-end models are included, as reported by Joopudi et al.^24^ (see the Discussion section). Highest values of the present study are highlighted in bold, with one result being outperformed by previous studies, denoted by an asterisk. Three different sources of benchmark data were utilized (here referred to as EU-ADR^53^, OMOP^54^ and FDA-rev^55^), from which a merged set was created, as the individual datasets had duplicate entries and varying sizes. Acc: accuracy; AUROC: area under the receiving operating characteristic curve; AUPRC: area under the precision–recall curve

Through node-based, hierarchical semantic similarity measures^52^, we can calculate the gain norm for the MedDRA^®^ terminology network as well. Comparison to the embedding-based gain norms (Fig. 5c) reveals a stark difference in the distinguishability of these validated groups from the background noise. While the embedding-based values consistently reach 0.25-0.3 across all measures (except Euclidean), going as high as 0.46, the MedDRA-based ones are all in the range of 0.02-0.04. Distributions plotted in Fig. 5a reveal that the MedDRA-based measures are hindered by the larger validated groups, which blend in with the random groups. An exception is the Jiang–Conrath measure that shows a clear separation, albeit with a very small margin. The low and sometimes even negative gain norm measured by the Euclidean distance in the case of the NSG model can be attributed to the use of dot product in its output layer (as opposed to the NTX model that uses cosine similarity instead), which resulted in much greater magnitudes and thus greater Euclidean distances among the embedding vectors. Given the well-known limitations of the Euclidean distance in high dimensions^49^, it becomes evident why information is much easier retrieved from the embedding space via the cosine similarity.

Overall, this evaluation demonstrates that even though the embedding vectors are organized into a different structure as compared to the MedDRA^®^ terminology network, drug-safety related information is better reflected by their emerging patterns, correlating to real life pharmacovigilance data, which could provide a more suitable resource in practice.

### 2.4. Causality prediction

As the focal point of our work, here we present an application of the embedding vectors as input features for a deep neural network model (Fig. 2d) performing drug–event causality prediction. The previously used validated drug–ADR datasets^46–48^ provide us only with positively labeled associations, which are insufficient for supervised training. Although there are datasets with negative pairs^53–55^, they are substantially smaller in scale and utilized mostly as benchmarks during performance evaluations. We followed this trend, resorting to synthetic negative data for training, which can be difficult to create, as easy negatives will lead to an overestimated model performance during training and validation, but poor generalization when tested on real samples. On the other hand, with hard negatives we might create an unsolvable classification task while also risking the inclusion of too many false negative examples. To this end, we defined six different negative sampling (i.e. random recombination) methods of various degrees of difficulties for the creation of the training set, using exclusively the drugs and reactions that are present in the validated drug–ADR data. Figure 6 shows the results of the same classifier trained, with different input embeddings, but with the same, best performing negative sampling approach, which coincides with the most difficult training set created out of the six. This clearly demonstrates that robust generalization is best achieved with hard negatives, when trivial, distinguishable patterns are avoided, forcing the model to exploit more complex underlying information. See Supplementary Table 10 and Supplementary Discussion 1 for further assessment on the performance-influencing effects of the various types of negative sampling approaches.

For a self-baseline comparison, we trained a version of the classifier that receives only the one-hot vector representation of reactions (“No embedding”) and learns the embeddings in-place, during training, so that the added benefit of using separately learnt reaction embeddings can be observed. The results of this simple ablation study (Fig. 6) clearly indicate one of the primary detriment of learning embeddings in-place, i.e. that many of the reactions in the benchmark datasets do not appear in the training data, making it impossible for the model to generalize to unseen samples, which can lead to a random classifier (AUROC ∼0.5) in practice. To position our embeddings in the larger scope of the modern representation learning landscape, we also trained versions of the same classifier using embedding vectors that are generated by various state of the art BERT-based models, specifically made for clinical and biological language tasks (PubMedBERT^31^, BioMed-RoBERTa^32^, Clinical ModernBERT^30^, BioClinical ModernBERT^29^). Although encoder-only LLMs are rarely employed to create static embeddings (as discussed in the Introduction section), their usage here demonstrates how natural-language-informed reaction embeddings compare to those trained on structured pharmacovigilance data. As the overall “Merged” results show (Fig. 6), the NSG-variant contrastive embeddings all outperformed the LLM-based ones, while the NTX-variants outperformed one of them. To reveal further insights about the structure of these LLM-based embeddings, we may employ the same dispersion analysis of related reactions as in Section 2.3, using a test for homogeneity of multivariate dispersions and the gain norm measure. Interestingly, only the reaction embeddings made by PubMedBERT show a significant difference of dispersions between validated and random groups, with quite a small margin (Supplementary Table 9), suggesting that LLM-based representation spaces do not capture interpretable drug-safety related information and rely on other patterns that are harder to explain. As the final baseline, we calculated the performance of the reporting odds ratio (ROR), which is a statistical disproportionality measure and the cornerstone of the signal detection method prominently used and recommended by EMA^56^. With only two exceptions (AUPRC for NTX-reaction2drug and NTX-reaction2all), all six contrastive embeddings resulted in the classifier outperforming the ROR method when evaluated on the merged benchmark set (Fig. 6), which demonstrates the ability of the embeddings to serve as a general resource in the support of decision making, improving the causality assessment process done by experts.

Lastly, we may compare the embeddings to each other. Overall, the reaction2reaction sampling shows better performance for the NTX model than both reaction2drug and reaction2all, matching their semantic evaluation trends (Fig. 4a). From this, we can conclude that reaction2reaction is the most suitable sampling method for the NTX model, possibly because it resembles the original use case of such architectures, which includes pairing together augmented versions of the same images to learn their embeddings. On the other hand, the reaction2all sampling method was not only the best performing NSG-variant but it also resulted in the overall best classification results across all methods (Fig. 6). We can observe a clear increase in the performance metrics going from reaction2drug to reaction2reaction to reaction2all for the NSG model, which might be explained by the available information (number of context terms) also increasing in this order. In general, embeddings made by NSG performed better than their NTX counterparts, which suggests that the N-pairs-based noise contrastive technique employed by NTX is less suited for learning effective embeddings from AERs as compared to the negative sampling noise contrastive technique of NSG.

## 3. Discussion

Here, we presented a framework to generate vector representations of adverse events from spontaneous AERs to offer an interpretable and effective resource for general machine learning applications in pharmacovigilance. A comprehensive analysis of the results was provided, revealing numerous mathematical and semantic patterns which imply high levels of preserved, drug-safety related information, with anatomical and pathophysiological patterns emerging.

To showcase a possible application of the embedding vectors and demonstrate their effectiveness as input features, we presented a deep neural classifier performing drug– event causality prediction. We trained the model on data that is independent of AERs used for the embeddings, as not to reuse them and to avoid some of the complications arising if we labeled parts of the reported pairs as positives and others as negatives via some mathematical or other unfunded consideration. An example with this issue would be the Transformer-based Hierarchy-aware Adverse Reaction Embeddings (T-HARE) model introduced by Joopudi et al.^24^, which is a type of representation learning we already discussed in the Introduction section, one that learns and uses the embeddings for a specific end-to-end prediction task instead of general purpose. For its training, a bipartite graph from FAERS data was constructed, and an edge-sampling method was utilized to select positive and negative pairs, based on their drug-normalized reporting frequency. Similar pharmacovigilance networks can indeed be used to perform signal detection, as shown by Pétervári et al.^57^ and Barbieri et al.^58^. However, the edge weights that are based on report counts need to be properly normalized via network topological features even for outperforming the ROR method^57^, suggesting that simpler normalization techniques might be inadequate for creating ground truth training data for machine learning. This limitation was indeed observed by Joopudi et al. during the training of T-HARE, which they remedied by scaling the loss on negative samples by their normalized pointwise mutual information. Another notable case is the aer2vec model introduced by Portanova et al.^17^ which is a similar negative sampling skip-gram version of word2vec as our NSG model, however, it was utilized to perform causality prediction directly, after training it on FAERS data. The first issue with this approach is that such shallow, noise contrastive embedding models are not meant to perfectly optimize their training objective, making them a suboptimal choice for this purpose. The other possible issue is the same as with T-HARE, that given its training regimen, the model might learn that all frequently reported pairs are positives, while only the rarely or never reported noisy pairs are negatives, which might fail to be of any use in practice. This problem of mishandling AER data for model training purposes in pharmacovigilance also extends to the creation of large drug–ADR association datasets as well, which often employ methodologically unsound assumptions, such as considering AERs with only a single reported suspect drug as being a confirmation of the causal link between that drug and the accompanying events^59,60^. Spontaneous AERs reflect subjective observations and opinions, which require comprehensive scientific evaluation involving multiple contextual factors, as explicitly stated by EMA in their official disclaimer (https://www.adrreports.eu/en/disclaimer.html). Using such AERs as confirmed positives on the same level as curated drug–ADR pairs extracted from public documents and package inserts^46–48^ can misrepresent the central problem of pharmacovigilance and result in a model with misleading performance metrics regarding its real life applicability.

This leads us to the most important limitation of studies dealing with machine-assisted causality assessment, one that plagues countless other biomedical research areas as well: the lack of proper gold standard benchmark data. Even with our diverse selection of sources (EU-ADR^53^, OMOP^54^, FDA-rev^55^), we faced the issue of having a very small test dataset, which imposes serious limitations on estimating real life performance for causality prediction models. As there are no comprehensive large-scale benchmark data with curated positive and negative drug–event associations, and many of the existing smaller sets are either unavailable^61^ or heavily restricted in scope^62,63^, we resorted to some of the more commonly used datasets, enabling comparison with results from other studies (Fig. 6). However, a closer look into these datasets reveals that they as well contain varying degrees of easy negatives, which is also reflected in the performance of the discussed methods. For example, in the EU-ADR dataset, negatives are defined as pairs without MEDLINE citations and AERs in VigiBase, making it the easiest dataset to predict, as also seen in the associated AUROC values (ROR: 0.9094, aer2vec: 0.9120, T-HARE: 0.9230, NSG-reaction2all: 0.9213). The OMOP dataset, on the other hand, was sourced via rigorous literature review and specifically aimed at providing a broad set of positive and negative controls. This is well reflected in the AUROC values (ROR: 0.7096, aer2vec: 0.8210, T-HARE: 0.8430, NSG-reaction2all: 0.9098) in which our model achieved a significant improvement compared to the other methods, potentially due to its training data being independent from AERs. Finally, in the FDA-rev dataset, negative controls were the result of a nearly identical negative sampling approach to what we used for generating the training data of the best performing classifier, aiming at a balanced distribution of drugs and reactions on both the positive and negative side while adhering to known associations. This led to the hardest negative samples and arguably the most difficult benchmark dataset, on which even our best model was slightly outperformed by the ROR method (AUROC: 0.7544 vs 0.7592). Low performance might also be attributed to the FDA-rev positive pairs containing not just validated reactions but “warnings and precautions” as well, which are significant safety hazards with or without established causality (see Supplementary Table 11 for our assessment). There are no performance measures available on FDA-rev data for the other methods.

In conclusion, we advance machine-assisted pharmacovigilance by offering a versatile and interpretable resource of adverse event vector representations that could enable the development of a wide array of machine learning applications to support decision making and improve patient safety.

## 4. Methods

### 4.1. Data sources

**Adverse event reports:** Raw AERs were accessed in XML format from the publicly available quarterly data extracts of the FAERS database^5^ for the interval 2012/Q4 – 2023/Q1, totaling more than 15M AERs. The beginning of the cutoff date corresponds to the modern FAERS database succeeding the legacy one (LAERS). Each AER contains a list of drugs and a list of adverse events (reactions), along with some constantly present information, like qualification of the reporter, seriousness, reporting date or reporting country, and some inconsistently available data, like patient sex, age or weight. Drugs have a “role” assigned by the reporter, indicating their possible connections to the case: suspect, interacting, concomitant. Additionally, the name of the active substances and indications (reason for prescription) may also be present. Reactions are listed as MedDRA^®^ PTs and may have an assigned “outcome”, like recovering or fatal.

**Drug names:** Given that drug names are not standardized in FAERS, multiple datasets were employed to identify the canonic DrugBank substance name of each drug, when possible, enabling us to trivially link them to their molecular attributes in DrugBank and PubChem. Our primary resource was RxNorm (monthly release 2023.07.03.)^64^, which is a collection of several drug vocabularies and a part of the Unified Medical Language System, maintained by the United States National Library of Medicine. We also utilized lists of synonyms and product names from DrugBank itself (full database, version 2023.01.04.)^11^, and the Anatomical, Therapeutic and Chemical (ATC) classification system (version 2021.12.03.)^65^, which is a hierarchical categorization of drugs (similar to how MedDRA^®^ is for reactions) maintained by the WHO Collaborating Centre for Drug Statistics Methodology. Although both DrugBank and ATC names are already present in RxNorm, we found that their separately sourced versions have additional entries.

**Reaction names:** MedDRA^®^, the Medical Dictionary for Regulatory Activities, is an extensive, international medical terminology for the support of regulatory communication and data analysis in pharmacovigilance, developed under the auspices of the International Council for Harmonisation of Technical Requirements for Pharmaceuticals for Human Use (ICH)^33^. MedDRA^®^ trademark is registered by ICH. Reactions in FAERS are already standardized to MedDRA^®^ PTs, with a corresponding MedDRA^®^ version specified for each reaction. However, MedDRA^®^ is updated on a biannual basis and access is tied to a paid license, thus we used only the most current version of the time (26.0).

**Drug features:** To provide molecular descriptors of drugs for the drug–event causality prediction model (Fig. 2d), a combination of DrugBank (full database, version 2023.01.04.)^11^ and PubChem (accessed on 2023.07.18.)^12^ data was used. While PubChem belongs to the United States National Library of Medicine, DrugBank is maintained by the Metabolomics Innovation Centre, Alberta, and the University of Alberta, Canada. We found that the chemical properties, such as “Rotatable bond count”, “Hydrogen bond acceptor count” or “Molecular weight”, are more consistently available on PubChem, and that several drug profiles on DrugBank also have an assigned “PubChem compound ID”, enabling the linking of the two data sources. Thus, from DrugBank, only the so-called biological properties of drugs were utilized, consisting of the major molecular biological determinants of their pharmacodynamics (targets) and pharmacokinetics (enzymes, carriers and transporters) properties^66^.

**Validated drug–ADR datasets:** Validated (positive only) drug–ADR associations were collected from three sources: SIDER^46^, IMI-PROTECT^47^ and BNF^48^. The Side Effect Resource (SIDER) is one of the most used datasets in pharmacovigilance research, featuring a large number of drug–ADR pairs that were text-mined from public documents and package inserts. However, its latest version (4.1 – 2015.10.21.), which was used for this study, is rather dated by now. The so-called “PROTECT” project of the Innovative Medicines Initiative (IMI) was conducted in a public–private partnership between the European Union (EU) and the European Federation for Pharmaceutical Industries and Associations. During its course, ADRs were collected from Section 4.8 of the Summary of Product Characteristics of medicinal products authorized in the EU. Our version of the IMI-PROTECT dataset (2017.06.30.) was accessed through the Internet Archive^67^, as the official link^68^ was broken at the time of writing. Finally, we utilized drug–ADR data which was collected from the British National Formulary (BNF) website^69^ by Kontsioti et al.^48^ (2018.06-08.).

**Benchmark sets:** Positive and negative drug–ADR associations were collected from three sources: EU-ADR^53^, OMOP^54^ and FDA-rev^55^. The EU-ADR project (Exploring and Understanding Adverse Drug Reactions by integrative mining of clinical records and biomedical knowledge) collected pairs for 10 selected reactions using MEDLINE citations and VigiBase reports, or the lack there of. The Observational Medical Outcomes Partnership (OMOP) project collected pairs for four selected reactions through systematic literature review and the use of product labels. Finally, Harpaz et al.^55^ utilized the MedWatch website of FDA to access drug label changes (revisions) as the source of positive pairs, and randomly recombined the same drugs and reactions with each other for negative pairs, after checking them against known associations (here referred to as FDA-rev).

### 4.2. Data pre-processing

**Report filtering:** We selected AERs only with the report type of “spontaneous”, which corresponds to those made by e.g. healthcare professionals, patients and pharmaceutical companies, but excludes studies. Mandatory AERs from companies may have multiple versions with the same case ID but different dates, corresponding to follow-up reports with added information. Consequently, only the most recent versions were kept, in-line with FAERS recommendations. These filtering steps resulted in about 10M reports remaining.

**Drug and reaction identification:** We only considered the reported drugs with the role of “suspect” which corresponds to the drugs primarily suspected to be the cause of the reactions in the given AER. To map the drugs into canonic DrugBank substance names, a combination of RxNorm, DrugBank and ATC data was utilized. Additionally, if the reported drug was specified as a composition of multiple substances, then the mapping was attempted with each substance, generating a new drug entry for each successful mapping. Reactions were mapped into MedDRA^®^ PTs through utilizing both the PTs and the lowest level terms (LLTs) of the most recent MedDRA^®^ dataset at the time. As a final AER filtering step, AERs that did not have at least one identified suspect drug or at least one identified reaction were excluded. This resulted in the exclusion of 197k reports, out of which only 35 were excluded because of unidentifiable reactions.

**Context-sampling for embedding:** AERs were reduced to simple lists of target–context pairs (Fig. 1a), where the target term is always a reaction and the associated context term is determined by the type of context-sampling: only drugs in the same AER (reaction2drug), only other reactions in the same AER (reaction2reaction), or both (reaction2all). Recurring pairs were kept to reflect their co-occurrence frequencies, and terms with a frequency of 10 or less were excluded to alleviate the problem of extremely rare tokens. Context sampling resulted in 14 765 target terms, 3908 context terms and 59M pairs for reaction2drug, 17 444 target/context terms and 174M pairs for reaction2reaction, and 18 130 target terms, 21 349 context terms and 233M pairs for reaction2all. The difference in term counts is caused by our frequency-based filtering condition.

**Drug feature collection:** The DrugBank database was filtered for entries that have a PubChem compound ID, enabling the linking of the two data sources. Following this filtering, 8724 drugs were kept, out of which 8345 had an actual “live” PubChem profile (i.e., status is not “non-live”). From DrugBank, the so-called biological features were utilized, i.e. lists of known targets, enzymes, carriers and transporters, that were filtered to include only proteins, and only those with a known “action” assigned to them. This resulted in a 2361-long multi-hot vector for each drug, indicating the presence or absence of these proteins. From PubChem, we restricted our selection to features that are available for all the selected drug substances, resulting in 18-long continuous-valued vectors.

**Classifier training/test data preparation:** The three validated drug–ADR datasets (SIDER^46^, IMI-PROTECT^47^, BNF^69^) and the three benchmark sets (EU-ADR^53^, OMOP^54^, FDA-rev^55^) were subjected to the same drug and reaction identification processes as the AERs, then merged separately, eliminating duplicate entries, to serve as the basis for a training and a test dataset, respectively. As a result, the merged validated drug–ADR dataset contained 1619 drugs, 4949 reactions and 165 260 pairs, which were also utilized for our dispersion analysis (Fig. 2c, Fig. 4). To best assess the performance of our drug– event causality classifier model on unseen cases, both positive and negative benchmark pairs were removed from the merged validated drug–ADR dataset for the training, before performing negative sampling. Furthermore, pairs from both the merged validated drug– ADR and benchmark sets had to be removed if they lacked drug features or reaction embedding vectors. For example, in the case of reaction2drug, which has the smallest reaction coverage, we had 1255 drugs, 4519 reactions and 138 162 pairs remaining. To increase the available test data, we manually identified the DrugBank canonic names and MedDRA^®^ PTs for the benchmark entries that were excluded by our automatic identification process. However, there still remained entries which had to be left out, such as drugs that refer to entire categories instead of individual substances. This resulted in 245 drugs, 46 reactions and 589 pairs remaining, with a 0.433–0.567 positive–negative ratio.

### 4.3. Embedding

Contrastive learning can be considered as a self-supervised approach, where the labels are not provided by ground truth data but rather determined by a given association, like the spatial proximity or semantic similarity of the samples^70^. Consequently, contrastive learning can be utilized for metric learning, a type of representation learning where the original associations between samples are preserved in the embedding vectors as a distance or similarity measure^71^. Both NTX and NSG are a form of noise-contrastive estimation (NCE), where the associated word pairs (positive label) are contrasted with noisy random pairs (negative label). A typical size for embedding vectors is between 100-300 dimensions, with vectors larger than 200 showing diminishing returns in quality^72^. Given the relatively small scale of our vocabularies (∼20 000) as compared to natural language corpora, we chose the dimension of 200 for the embedding vectors, same as what several other medical and biological static embeddings use^19,73^.

**NTX:** The term “normalized temperature-scaled cross entropy loss” (NT-xent) was coined by Chen et al.^36^ in their work where they described a simple framework for contrastive learning of visual representations (SimCLR), but the formulation of the loss itself is almost identical to the N-pairs loss^74^ and the loss of InfoNCE^75^. In all cases, noise pairs are not sampled explicitly but rather derived from the associated pairs themselves. In each batch of *N* associated pairs, we can define *N*(*N* − 1) non-associated ones by recombination. This leads to a multiclass classification problem for both constituents (*t*: target, *c*: context) of the *N* associated pairs, with the overall loss function defined as:

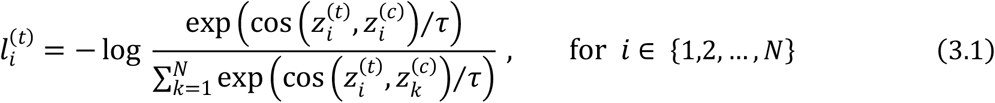

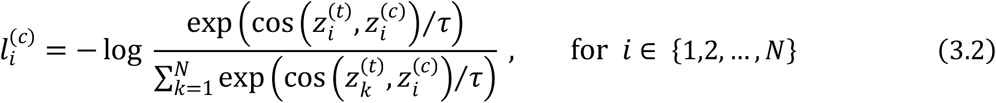

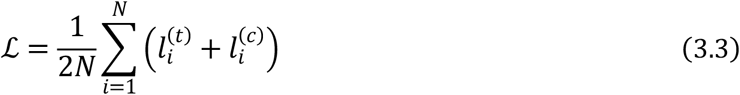

where cos(⋅,⋅) refers to the cosine similarity (i.e. normalized dot product), *τ* is the temperature parameter, and 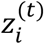 and 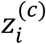 are the vector representations of the target and context terms, respectively, after a shared, two-layer projection head:

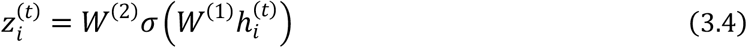

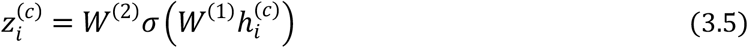

with 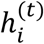 and 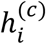 being the embedding vectors that are kept for downstream tasks, and the projection head consisting of two fully connected layers (*d* = 128) with a rectified linear unit (ReLU) activation function in-between.

Given that the model optimizes for all other *N*(*N* − 1) pairs in the batch to be as dissimilar as possible, we constructed the batches with no duplicate terms in order to avoid a controversial optimization task. This was achieved by dividing the list of unique target terms to batch-sized groups, then to each target we assigned one randomly sampled context from its corresponding list of contexts, resampling them until all contexts are unique inside the given batch. We considered the resulting batches of target–context pairs to be data for a single epoch, and generated a new set for each epoch. During sampling a context term for a given target term, it was not checked whether it is a context for the other targets in the batch as well, only whether it was already in the batch or not. This is in line with the assumption of NCE that the influence of accidental hits will be diminished by sufficient amounts of data. On a side note, it is possible that our NTX model would perform better if the selection of pairs resembled more closely the augmentation of images, as done in SimCLR, such as by randomly swapping the reactions for one of their semantically closest neighbors in a given terminology. This would obviously deviate from our original aim of utilizing AER data and therefore, it is left as a future direction to be explored.

Training was done for 1000 epochs (embeddings extracted after every 100^th^), using Root Mean Square Propagation (RMSProp) with learning rate *η* = 0.01, temperature parameter *τ* = 0.3, and a batch size of 16.

**NSG:** Accompanying the skip-gram word2vec model^76^, Mikolov et al. introduced several extensions in their concurrent work, namely the use of subsampling and negative sampling^37^, both of which were utilized in our NSG model. Negative sampling is an alternative to hierarchical softmax, transforming the word2vec objective from a vector-to-vector prediction task into a binary classification of vector pairs, while significantly reducing the required computations. The core idea is that rather than considering all but the currently associated term as negatives in each pass (which requires tuning all weights at the same time), only a handful of negatives are used, sampled randomly from a noise distribution *P*_*n*_(*c*). Hence, the loss function is given by:

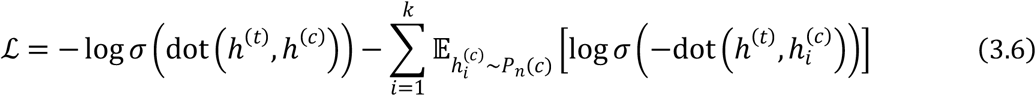

where dot(⋅,⋅) refers to the dot product, *σ* is a sigmoid activation function, *h*^(*t*)^ and *h*^(*c*)^ are the embedding vectors of an associated target–context pair, respectively, and 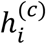 are the embedding vectors of the sampled negatives, which are not required to be unique, only to not match the current associated context. We set *P*_*n*_(*c*) to be the smoothed unigram distribution:

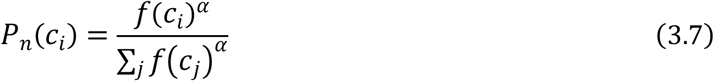

where *f* is the number of occurrences of the given context term in the report corpus and *α* is the smoothing parameter. Training data was constructed by going through all associated target–context pairs of the corpus and assessing the chance of the target term being kept, according to the subsampling distribution:

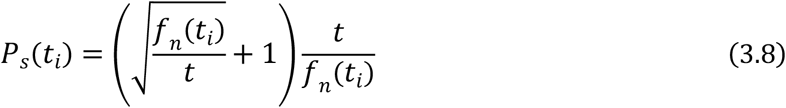

where *f*_*n*_ is the normalized occurrence count of the given target term and *t* is a hyperparameter governing the “aggressiveness” of the subsampling, i.e. how likely it is that a target term above a certain count will be discarded. This alleviates the severity of imbalance observed in the occurrences of terms, ranging from 11 (the minimum threshold set during pre-processing) to 1-3M. Negative sampling was done after subsampling, selecting *k* contexts as negatives to each kept target. The resulting data from one pass through the entire corpus was considered as training data for one epoch, with training data being regenerated after every 5^th^ epoch, starting from the subsampling step.

Training was done for 30 epochs (embeddings extracted after every 5^th^), using RMSProp with learning rate *η* = 0.01, smoothing parameter *α* = 0.75, subsampling parameter *t* = 0.00001, number of sampled negatives *k* = 7, and 8 positive target–context pairs per batch, thus a total batch size of 64.

**LLMs:** Reaction embeddings were created via the various pretrained BERT-based models^29–32^ by providing them as input the same list of reaction terms as used for the reaction2all embeddings. Each term was embedded separately, like a sentence, and mean pooling was applied to generate single static embeddings, which is a method known as decontextualization^34,35^.

### 4.4. Vector similarity measures

To determine the embedding distances for *t*-SNE (Fig. 3) and for the semantic evaluation of the top 10 closest neighbors (Fig. 4), we utilized cosine distance (i.e. 1 − cosine similarity) and cosine similarity, respectively. Cosine similarity is the most common vector-similarity measure for word vectors because the embedding models producing them also include either dot product or its normalized version (i.e. cosine similarity) in the output layer, before non-linearity. As described in Section 4.3, the final output value is either maximized for associated word pairs or minimized for non-associated ones, thus resulting in the associations being preserved by the used measure (metric learning).

For the dispersion analysis (Fig. 5), we used cosine similarity, dot product, and three more vector-similarity measures of interest: (i) the Euclidean distance, as the embedding spaces are technically Euclidean, even though the shape of the manifolds and their high dimensionality make the Euclidean distance here less applicable, (ii) the Bray–Curtis dissimilarity (also called Bray–Curtis distance or Sørensen dissimilarity/distance) and (iii) the Canberra distance, both of which are weighted versions of the Manhattan distance, being prominently used in ecology and clustering-based classification methods^77–79^. All three of these measures are negated in our dispersion calculations (Fig. 5a), so that higher values correspond to closer associations across all methods.

### 4.5. Hierarchical semantic similarity measures

It is important to mention that MedDRA^®^ is not a taxonomy, as individual PTs and higher level terms can belong to multiple parent categories, resulting in several possible intersecting “paths” through which the highest level (SOC) can be reached from a PT. There is, however, always a path denoted as the primary one, which we utilized in our clustering analysis (Fig. 3c). For the dispersion analysis (Fig. 5), we considered four MedDRA-based similarity measures^52^, namely: (i) mean minimum path, which is the inverse distance between the two reaction PTs, averaged across all of their possible paths, or equivalently, the inverse sum of their distances from their lowest common ancestors (LCAs), averaged across all of their possible paths, (ii) Resnik similarity, which uses the average information content of the LCAs from all paths, where information content is calculated using the report corpus, (iii) Jiang–Conrath and (iv) Lin similarities, both of which are normalized versions of the Resnik similarity. An “imaginary” root node was inserted above the SOC terms, connecting all of them, so that these calculations are still possible for PTs that otherwise do not have an LCA.

### 4.6. Visualization and clustering

**Dimensionality reduction:** *t*-SNE optimizes the composition of local neighborhoods in the target space (*n*_*components* = 2 here), best preserving how they appeared in the source, while global placements are often arbitrary^39^. Furthermore, small patterns might emerge that are mere artifacts of the optimization process. Larger, well-separated clusters, however, are still consistently retrieved^80^, which provides sufficient insight in our case. Neighborhood size in the source dimension is governed by the perplexity parameter, determining the number of closest data points for each embedding vector to be considered. Closeness is defined by a given distance or vector-similarity measure, such as Euclidean or cosine. We used the scikit-learn^81^ implementation of *t*-SNE with *perplexity* = 30, *metric* = “*cosine*”, a maximum number of iterations *n*_*iter* = 3000, and *random*_*state* = 42, for reproducible results. Other parameters were left as default.

**Clustering:** HDBSCAN is the hierarchical adaptation of the Density-Based Spatial Clustering of Applications with Noise (DBSCAN)^40^, which groups points into a cluster if they are in the proximity of at least a minimum number of other points, in a set distance, determined by the epsilon parameter. DBSCAN performs clustering iteratively, thus if there is a cluster in epsilon distance to a yet-to-be clustered point, that new point is added to the existing cluster. HDBSCAN modifies DBSCAN by running the algorithm multiple times with different epsilons, detecting clusters at variable densities, and merging the most stable ones as its final result. We used the scikit-learn^81^ implementation of HDBSCAN, empirically setting *min*_*cluster*_*size* = 10, while the other parameters were left as default, most importantly *min*_*samples* = *min*_*cluster*_*size* and *metric* = “*euclidean*”.

### 4.7. Test for homogeneity of multivariate dispersions

Anderson^50^ proposed a permutation test for homogeneity of multivariate dispersions (“permdisp”) that enables the use of arbitrary distance measures, and calculates group dispersion by considering the distances of samples from the estimated group center. As described by Gijbels and Omelka^49^, the test can be modified (“modified permdisp”) to use pairwise interpoint distances instead. This version of the test was not only used directly (Supplementary Table 9) but it also served as an inspiration for the “gain norm” dispersion difference measure (Section 4.8), which can be clearly seen from their formulations.

The test statistic *F* for the modified permdisp can be calculated between any *K* number of groups by:

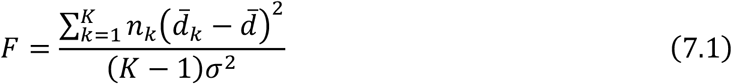

where *n*_*k*_ is the size of the k^th^ group with 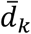 being the average of its pairwise interpoint distances, and 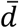 being the global average of the entire distance matrix. The asymptotic variance *σ*^2^ is estimated by the jackknife estimator:

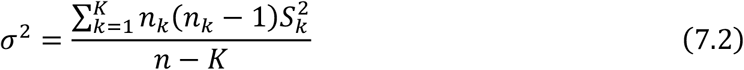

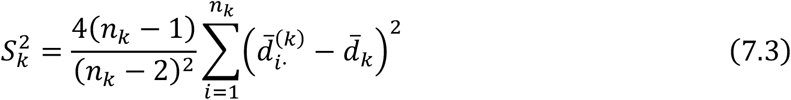

where 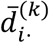 is the average of the elements belonging to the k^th^ group in the i^th^ row of the distance matrix, without the diagonal. This is equivalent to taking the average of the distances from data point *i* in group *k* to every other data point in that group.

To obtain a p-value, the distribution of the test statistic can be estimated by a permutation procedure, where the assigned group labels of the data points are repeatedly shuffled by sampling from all permutations. We also implemented improvements to the properties of the permutation procedure, as described by Gijbels and Omelka^49^, that can be achieved through performing Principal Coordinate Analysis (PCoA) transformation on the distance matrix in order to reduce the distances among groups. For this step, it is best to avoid negative eigenvalues, which can be done by applying the Cailliez correction method on the distance matrix (Supplementary Note)^79^.

For each validated group of reactions, we created a randomly sampled group of the same size, and performed the modified permdisp test between the two (*K* = 2) with a maximum permutation sampling count of 999. Due to multiple testing (all validated–random group pairs), the resulting permutation *p*-values were corrected by the Benjamini–Hochberg procedure^82^, yielding *q*-values. Results are summarized for each embedding, with each measure, as the ratio of validated reaction groups showing a significantly (*q* < 0.05) different dispersion compared to the corresponding random reaction groups (Supplementary Table 9).

### 4.8. Gain norm

The use of another dispersion difference measure^51^, here simply termed “gain norm”, was inspired by the previously described modified permdisp test, aiming at an intuitive and finer way of describing how well the validated groups of reactions separate from noise in the embedding space. Calculation involves the difference of [0,1]-normalized average pairwise interpoint distances:

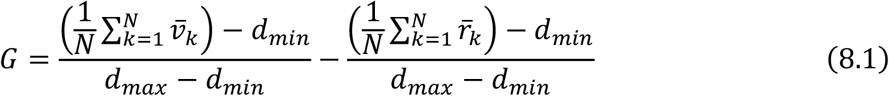

where *N* is the number of validated groups and, consequently, the number of same-sized random groups, with 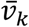 and 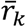 being the average pairwise distances within the k^th^ validated and k^th^ random groups, respectively, and *d*_*min*_ and *d*_*max*_ being the global minimum and maximum measured values. With large enough group sizes and high enough *N*, the average of the random groups 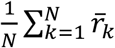 approximates the global average 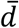, thus we can rewrite the formula so that it resembles the test statistic from Eq. 7.1 a little closer:

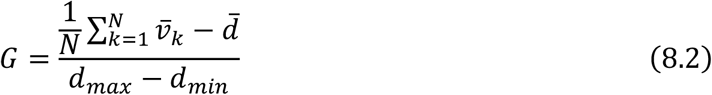

In brief, the test statistic *F* for the modified permdisp test calculates the sum of squared differences between the average pairwise distances of each group and the global average, scaling them with group sizes, and divides the result with the asymptotic variance *σ*^2^ (Eq. 7.1-7.3). Here, we calculate the average difference between the average pairwise distances of each group and the global average, dividing it by the complete range of the measures for normalization (Eq. 8.2).

### 4.9. Generating negative training data

Starting from a list of validated (positive) drug–ADR pairs, two aspects were considered for the generation of negative pairs: which constituent of the positive pairs to keep (drug, reaction, none) and whether to sample the other constituent uniformly from the available terms or follow their frequency distribution as observed in the positive pairs. This resulted in six negative sampling methods, out of which the one where we keep the reaction and sample drugs with their frequency distribution proved to be the best performing one (see Supplementary Table 10 and Supplementary Discussion 1). In all cases, the same number of negative pairs were created as there are positives, with the only criterion of not matching any of the existing positives. Drug and reaction terms available for sampling were restricted to those that are present in the validated data. Additionally, oversampling was performed by repeating this generation step 5 more times, combining the resulting data with the positive pairs for each repetition, thus the final training data was 12 times the size of the original validated drug–ADR data.

### 4.10. Drug–event causality prediction model

With this prediction model, our aim was to construct a relatively simple deep neural classifier to demonstrate how our embeddings can be utilized for drug–event causality prediction. Therefore, the input of this model consists of the embedding vector of the given reaction as well as the chemical and biological feature vectors of the given drug. These drug feature vectors are separately pre-processed by a dense layer each (*d* = 100), without an activation function, bringing them in-line with the reaction embedding (*d* = 200) in terms of dimension, so that their contributions are more balanced. Following concatenation, the new vector (*d* = 400) is passed through two hidden layers (*d* = 1024 and *d* = 256), outfitted with batch normalizations and a drop out (0.4), as seen in Fig. 2d, with leaky ReLU activation functions in-between. The output, that is given by sigmoid activation and optimized against the expected label through binary cross entropy loss, can be considered as the probability of the given drug and reaction being causally associated.

Hyperparameter search was done by using the merged validated drug–ADR dataset, repeatedly splitting it to a training and test set (0.7/0.3) for each run. Negative data generation was performed separately on the two sets and in a way that the newly sampled negatives in the training set are not checked against the test set, whether they are false negatives or not, avoiding a possible data leak. For the final training, the entire validated drug–ADR data was employed, only filtering out those pairs that also appear in the benchmark sets. Then, with each of the six embeddings and with each of the six negative sampling methods, model training was performed 10 times, generating new negative data for each run (Supplementary Table 10).

Training was done for 50 epochs, using stochastic gradient descent with learning rate *η* = 0.001 and a batch size of 512.

### 4.11. Reporting odds ratio

The reporting odds ratio (ROR) is commonly used as the basis of signal detection in pharmacovigilance^56^. It calculates the statistical disproportionality between the probability of observing a given reaction in the presence of a given medicinal product and the probability of observing the same reaction in the absence of that product. Correspondingly, the ROR is given by the ratio of these odds:

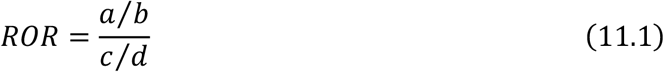

where *a* is the number of AERs containing both the given drug and the given reaction. Values *b* and *c* are the number of AERs containing only one of the constituents, either the given drug or the given reaction, respectively. These values correspond to their occurrences in other scenarios that may reflect how specifically or widely they are reported. Finally, *d* is simply the number of AERs without either of the constituents. We estimate the standard deviation *s* of ROR as:

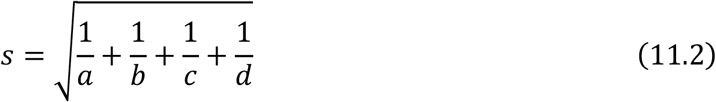

From this, the lower bound of the 95% confidence interval of the ROR value is calculated as:

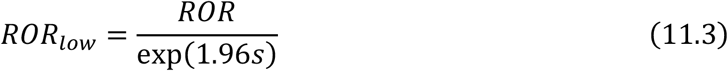

According to EMA guidelines^56^, when this value is equal to or greater than 1 and there are at least three reported cases (*a* ≥ 3), the given drug–event pair is considered to be a signal, meaning that their potential causal relation requires further investigations. For the performance evaluation of our classifier, the *ROR*_*low*_ values were calculated on the entire pre-processed report corpus, normalized to [0,1] and set to 0 where *a* < 3. This final step involved also the pairs that were never reported (i.e. *a* = 0), such as in the case of many of the negative entries in the benchmark sets, which heavily boosted the performance of the ROR method.

### 4.12. Software and hardware

Data pre-processing was implemented in C++ 17 standard, while model trainings and result analyses were done using Python (version 3.10). The contrastive embedding and classifier models were implemented using Tensorflow/Keras (version 2.11)^83^. We used the scikit-learn (version 1.3)^81^ implementations of *t*-SNE and HDBSCAN for dimensionality reduction and clustering analysis, respectively, and plotly (version 5.19)^84^ for visualization. BERT-based pretrained LLMs were accessed from Hugging Face through the *transformers* package (version 4.48)^85^, and used to generate embeddings (inference only) with Python (version 3.12) and PyTorch (version 2.6)^86^. Model trainings were performed on an NVIDIA A100 Tensor Core GPU, taking 1-3 hours for the NTX models and 46-221 hours for the NSG models, depending on the context-sampling method used. This marks a notable advantage in our NTX model as compared to the NSG model, in terms of efficiency. Classifier training took 1-2 hours per run, depending on the negative sampling methods used.

## 5. Data availability

All the data files for training our embedding models and the drug–event classifier, as well as for performing clustering are available at deepvigilace.com, namely the pre-processed report corpus made from FAERS adverse event reports, the derived drug and reaction frequencies, the pre-processed validated drug–ADR and benchmark data, the collected drug feature vectors, the presented reaction embedding vectors, and their two-dimensional *t*-SNE coordinates. In addition, the complete results of the HDBSCAN clustering, the raw top-neighbor tables for *Myocardial Ischeamia*, and the prediction results made by the ROR-based signal detection method and our best classifier on the benchmark sets were also made available, along with the best performing classifier model itself. Interactive HTML visualizations are provided for the *t*-SNE and clustering results, enabling manual data analysis and browsing. Due to licensing, MedDRA^®^ terminology data is not provided, which is needed for top 10 closest neighbor and dispersion calculations with the hierarchical semantic similarity measures.

## 6. Code availability

All code necessary for replicating the results of this study from the provided pre-processed adverse event reporting data, i.e. Python code for generating, analyzing and using the reaction embeddings, is available under the MIT license at github.com/semmelweis-pharmacology/deepvigilace.

## 7. Acknowledgements

Project No. RRF-2.3.1-21-2022-00003 has been implemented with the support provided by the European Commission, European Union. This project has received funding from the HUN-REN Hungarian Research Network and the National Research, Development and Innovation Office of Hungary: 2020-1.1.6-JÖVŐ-2021-00013. The research was financed by the Thematic Excellence Programme (2020-4.1.1.-TKP2020) of the Ministry for Innovation and Technology in Hungary, within the framework of the Therapeutic Development and Bioimaging thematic programmes of the Semmelweis University. This study has been supported by the Semmelweis 250+ Excellence Fellowship (O.M.B., E.P.). O.M.B was supported by the EKÖP-2024-23 New National Excellence Program of the Ministry for Culture and Innovation from the source of the National Research, Development and Innovation Fund. We thank Dóra Bihary for providing valuable feedback about the grammatical correctness and language of the manuscript.

## 8. Contributions

O.M.B.: conceptualization, methodology design, coding, model implementation, data analysis, visualization, paper preparation – original draft. M.P.: conceptualization, data analysis, domain expertise, paper preparation – revision. Á.M.Cs.: coding, data analysis, domain expertise, visualization, paper preparation – original draft. E.P.: data analysis, domain expertise, paper preparation – revision. A.H.: methodology design, paper preparation – revision. P.F.: project supervision, domain expertise, paper preparation – revision. B.Á.: project supervision, conceptualization, methodology design, paper preparation – revision.

## 9. Competing interests

P.F. is the founder and CEO of, and B.Á. is employed by, Pharmahungary Group, a group of R&D companies. M.P. is the founder and CEO of Sanovigado Kft, a pharmaceutical consultancy and R&D company.

## Notes

https://www.deepvigilace.com/

https://github.com/semmelweis-pharmacology/deepvigilace

### Summary of Updates

To position our embeddings in the modern representation learning field, new evaluations were conducted using embeddings generated by various state of the art BERT models. The narrative of the study was extended to include these results, improving our claims and clarifying the primary aim. Ablation study was added, demonstrating how the drug-event causality classifier performs without embeddings. Further assessment was added about the available pharmacovigilance data for training and testing machine learning models. Each results section was extended to provide some concluding statements.

## References

1. European Medicines Agency. Guideline on Good Pharmacovigilance Practices (GVP) Annex I-Definitions (Rev 4). https://www.ema.europa.eu (2017).

2. EudraVigilance | European Medicines Agency (EMA). https://www.ema.europa.eu/en/human-regulatory-overview/research-development/pharmacovigilance-research-development/eudravigilance.

3. FDA Adverse Event Reporting System (FAERS) Database | FDA. https://www.fda.gov/drugs/drug-approvals-and-databases/fda-adverse-event-reporting-system-faers-database.

4. Lindquist, M. VigiBase, the WHO Global ICSR Database System: Basic facts. Drug Inf. J. 42, 409–419 (2008).

5. FDA Adverse Events Reporting System (FAERS) Public Dashboard - FDA Adverse Events Reporting System (FAERS) Public Dashboard | Sheet - Qlik Sense. https://fis.fda.gov/sense/app/95239e26-e0be-42d9-a960-9a5f7f1c25ee/sheet/7a47a261-d58b-4203-a8aa-6d3021737452/state/analysis.

6. Danysz, K. et al. Artificial Intelligence and the Future of the Drug Safety Professional. Drug Saf. 42, 491–497 (2019).

7. Al-Azzawi, F., Mahmoud, I., Haguinet, F., Bate, A. & Sessa, M. Developing an Artificial Intelligence-Guided Signal Detection in the Food and Drug Administration Adverse Event Reporting System (FAERS): A Proof-of-Concept Study Using Galcanezumab and Simulated Data. Drug Saf. 46, 743–751 (2023).

8. Cherkas, Y., Ide, J. & van Stekelenborg, J. Leveraging Machine Learning to Facilitate Individual Case Causality Assessment of Adverse Drug Reactions. Drug Saf. 45, 571–582 (2022).

9. De Abreu Ferreira, R. et al. A Pilot, Predictive Surveillance Model in Pharmacovigilance Using Machine Learning Approaches. Adv. Ther. 41, 2435– 2445 (2024).

10. Bae, J. H. et al. Machine Learning for Detection of Safety Signals From Spontaneous Reporting System Data: Example of Nivolumab and Docetaxel. Front. Pharmacol. 11, 602365 (2021).

11. Wishart, D. S. et al. DrugBank 5.0: A major update to the DrugBank database for 2018. Nucleic Acids Res. 46, D1074–D1082 (2018).

12. Kim, S. & Bolton, E. E. PubChem: A Large-Scale Public Chemical Database for Drug Discovery. in Open Access Databases and Datasets for Drug Discovery 41– 66 (John Wiley & Sons, Ltd, 2023). doi:10.1002/9783527830497.ch2.

13. Mower, J., Subramanian, D. & Cohen, T. Learning predictive models of drug side-effect relationships from distributed representations of literature-derived semantic predications. J. Am. Med. Informatics Assoc. 25, 1339–1350 (2018).

14. Gattepaille, L. M. Using the WHO database of spontaneous reports to build joint vector representations of drugs and adverse drug reactions, a promising avenue for pharmacovigilance. in 2019 IEEE International Conference on Healthcare Informatics (ICHI) (Institute of Electrical and Electronics Engineers Inc., 2019). doi:10.1109/ICHI.2019.8904551.

15. Beam, A. L. et al. Clinical concept embeddings learned from massive sources of multimodal medical data. in Pacific Symposium on Biocomputing vol. 25 295–306 (World Scientific Publishing Co. Pte Ltd, 2020).

16. Choi, Y., Chiu, C.Y.-I. & Sontag, D. Learning Low-Dimensional Representations of Medical Concepts. in AMIA Summits on Translational Science Proceedings vol. 2016 41–50 (2016).

17. Portanova, J., Murray, N., Mower, J., Subramanian, D. & Cohen, T. aer2vec: Distributed Representations of Adverse Event Reporting System Data as a Means to Identify Drug/Side-Effect Associations. in AMIA Annual Symposium Proceedings vol. 2019 717–726 (American Medical Informatics Association, 2019).

18. Nikfarjam, A., Sarker, A., O’Connor, K., Ginn, R. & Gonzalez, G. Pharmacovigilance from social media: mining adverse drug reaction mentions using sequence labeling with word embedding cluster features. J. Am. Med. Inform. Assoc. 22, 671 (2015).

19. Chen, Q. et al. BioConceptVec: Creating and evaluating literature-based biomedical concept embeddings on a large scale. PLOS Comput. Biol. 16, e1007617 (2020).

20. Erlanson, N., China, J. F., Taavola, H. & Norén, G. N. Clinical Relatedness and Stability of vigiVec Semantic Vector Representations of Adverse Events and Drugs in Pharmacovigilance. Drug Saf. 48, 401 (2025).

21. Zhang, Y., Chen, Q., Yang, Z., Lin, H. & Lu, Z. BioWordVec, improving biomedical word embeddings with subword information and MeSH. Sci. Data 6, 52 (2019).

22. Zhao, H. et al. Identifying the serious clinical outcomes of adverse reactions to drugs by a multi-task deep learning framework. Commun. Biol. 6, 1–14 (2023).

23. Zhang, F., Sun, B., Diao, X., Zhao, W. & Shu, T. Prediction of adverse drug reactions based on knowledge graph embedding. BMC Med. Inform. Decis. Mak. 21, 1–11 (2021).

24. Joopudi, V., Dandala, B.Tsou, C.-H. & Liang, J. J. Hierarchy-aware Adverse Reaction Embeddings for Signal Detection. in AMIA Annual Symposium Proceedings vol. 2022 596 (2022).

25. Mao, Y. & Fung, K. W. Use of word and graph embedding to measure semantic relatedness between unified medical language system concepts. J. Am. Med. Informatics Assoc. 27, 1538–1546 (2020).

26. Alsentzer, E. et al. Publicly Available Clinical BERT Embeddings. in Proceedings of the 2nd Clinical Natural Language Processing Workshop 72–78 (Association for Computational Linguistics, Minneapolis, Minnesota, USA, 2019). doi:10.18653/v1/W19-1909.

27. Park, J. et al. Validation of a Natural Language Machine Learning Model for Safety Literature Surveillance. Drug Saf. 47, 71–80 (2024).

28. Portelli, B., Lenzi, E., Chersoni, E., Serra, G. & Santus, E. BERT Prescriptions to avoid unwanted headaches: A comparison of transformer architectures for adverse drug event detection. in EACL 2021 - 16th Conference of the European Chapter of the Association for Computational Linguistics, Proceedings of the Conference 1740–1747 (Association for Computational Linguistics (ACL), 2021). doi:10.18653/v1/2021.eacl-main.149.

29. Sounack, T. et al. BioClinical ModernBERT: A State-of-the-Art Long-Context Encoder for Biomedical and Clinical NLP. (2025).

30. Lee, S. A., Wu, A. & Chiang, J. N. Clinical ModernBERT: An efficient and long context encoder for biomedical text. (2025).

31. Gu, Y. et al. Domain-Specific Language Model Pretraining for Biomedical Natural Language Processing. ACM Trans. Comput. Healthc. 3, 24 (2021).

32. Gururangan, S. et al. Don’t Stop Pretraining: Adapt Language Models to Domains and Tasks. Proc. Annu. Meet. Assoc. Comput. Linguist. 8342–8360 (2020) doi:10.18653/v1/2020.acl-main.740.

33. Mozzicato, P. Meddra: An overview of the medical dictionary for regulatory activities. Pharmaceutical Medicine vol. 23 65–75 at 10.1007/BF03256752 (2009).

34. Bommasani, R., Davis, K. & Cardie, C. Interpreting Pretrained Contextualized Representations via Reductions to Static Embeddings. Proc. Annu. Meet. Assoc. Comput. Linguist. 4758–4781 (2020) doi:10.18653/V1/2020.ACL-MAIN.431.

35. Gupta, P. & Jaggi, M. Obtaining Better Static Word Embeddings Using Contextual Embedding Models. ACL-IJCNLP 2021 - 59th Annu. Meet. Assoc. Comput. Linguist. 11th Int. Jt. Conf. Nat. Lang. Process. Proc. Conf. 1, 5241–5253 (2021).

36. Chen, T., Kornblith, S., Norouzi, M. & Hinton, G. A Simple Framework for Contrastive Learning of Visual Representations. in Proceedings of the 37th International Conference on Machine Learning (eds. Iii, H.D. & Singh, A.) vol. 119 1597–1607 (PMLR, 2020).

37. Mikolov, T., Sutskever, I., Chen, K., Corrado, G. & Dean, J. Distributed representations of words and phrases and their compositionality. in Advances in Neural Information Processing Systems (Neural information processing systems foundation, 2013).

38. Gutmann, M. & Hyvärinen, A. Noise-contrastive estimation: A new estimation principle for unnormalized statistical models. in Journal of Machine Learning Research vol. 9 297–304 (JMLR Workshop and Conference Proceedings, 2010).

39. van der Maaten, L. & Hinton, G. Visualizing Data using t-SNE. J. Mach. Learn. Res. 9, 2579–2605 (2008).

40. Campello, R. J. G. B., Moulavi, D. & Sander, J. Density-Based Clustering Based on Hierarchical Density Estimates. in Advances in Knowledge Discovery and Data Mining (eds. Pei, J., Tseng, V. S., Cao, L., Motoda, H. & Xu, G.) 160–172 (Springer Berlin Heidelberg, Berlin, Heidelberg, 2013).

41. Roth, G. A. et al. Global, Regional, and National Burden of Cardiovascular Diseases for 10 Causes, 1990 to 2015. J. Am. Coll. Cardiol. 70, 1–25 (2017).

42. Rezende, P. C., Ribas, F. F., Serrano, C. V. & Hueb, W. Clinical significance of chronic myocardial ischemia in coronary artery disease patients. J. Thorac. Dis. 11, 1005 (2019).

43. Shimokawa, H. & Yasuda, S. Myocardial ischemia: Current concepts and future perspectives. J. Cardiol. 52, 67–78 (2008).

44. Szabados, T. et al. Identification of New, Translatable ProtectomiRs against Myocardial Ischemia/Reperfusion Injury and Oxidative Stress: The Role of MMP/Biglycan Signaling Pathways. Antioxidants 13, 674 (2024).

45. Nagy, R. N. et al. Cardioprotective microRNAs (protectomiRs) in a pig model of acute myocardial infarction and cardioprotection by ischaemic conditioning: MiR-450a. Br. J. Pharmacol. 182, 396–416 (2024).

46. Kuhn, M., Letunic, I., Jensen, L. J. & Bork, P. The SIDER database of drugs and side effects. Nucleic Acids Res. 44, D1075 (2016).

47. Wisniewski, A. F. Z. et al. Good Signal Detection Practices: Evidence from IMI PROTECT. Drug Saf. 39, 469–490 (2016).

48. Kontsioti, E., Maskell, S., Dutta, B. & Pirmohamed, M. A reference set of clinically relevant adverse drug-drug interactions. Sci. Data 9, 1–9 (2022).

49. Gijbels, I. & Omelka, M. Testing for Homogeneity of Multivariate Dispersions Using Dissimilarity Measures. Biometrics 69, 137–145 (2013).

50. Anderson, M. J. Distance-based tests for homogeneity of multivariate dispersions. Biometrics 62, 245–253 (2006).

51. Spector, J. et al. Transformers Enhance the Predictive Power of Network Medicine. medRxiv 2025.01.27.25321204 (2025) doi:10.1101/2025.01.27.25321204.

52. Bill, R. W. et al. Evaluating semantic relatedness and similarity measures with Standardized MedDRA Queries. in AMIA Annual Symposium Proceedings vol. 2012 43–50 (American Medical Informatics Association, 2012).

53. Coloma, P. M. et al. A reference standard for evaluation of methods for drug safety signal detection using electronic healthcare record databases. Drug Saf. 36, 13–23 (2013).

54. Ryan, P. B. et al. Defining a reference set to support methodological research in drug safety. Drug Saf. 36, 33–47 (2013).

55. Harpaz, R. et al. A time-indexed reference standard of adverse drug reactions. Sci. Data 1, 1–10 (2014).

56. European Medicines Agency. Screening for adverse reactions in EudraVigilance. https://www.ema.europa.eu/en/documents/other/screening-adverse-reactions-eudravigilance_en.pdf (2016).

57. Pétervári, M. et al. Network Analysis for Signal Detection in Spontaneous Adverse Event Reporting Database: Application of Network Weighting Normalization to Characterize Cardiovascular Drug Safety. Drug Saf. 45, 1423–1438 (2022).

58. Barbieri, M. A. et al. Network Analysis and Machine Learning for Signal Detection and Prioritization Using Electronic Healthcare Records and Administrative Databases: A Proof of Concept in Drug-Induced Acute Myocardial Infarction. Drug Saf. 1–14 (2025) doi:10.1007/S40264-025-01515-Y/FIGURES/4.

59. Yue, Q. X. et al. Mining Real-World Big Data to Characterize Adverse Drug Reaction Quantitatively: Mixed Methods Study. J. Med. Internet Res. 26, e48572 (2024).

60. Yu, Z., Wu, Z., Li, W., Liu, G. & Tang, Y. MetaADEDB 2.0: a comprehensive database on adverse drug events. Bioinformatics 37, 2221–2222 (2021).

61. Caster, O. et al. Assessment of the Utility of Social Media for Broad-Ranging Statistical Signal Detection in Pharmacovigilance: Results from the WEB-RADR Project. Drug Saf. 41, 1355 (2018).

62. Osokogu, O. U. et al. Pediatric Drug Safety Signal Detection: A New Drug–Event Reference Set for Performance Testing of Data-Mining Methods and Systems. Drug Saf. 38, 207 (2015).

63. Ito, S. & Narukawa, M. Development of a Drug Safety Signal Detection Reference Set Using Japanese Safety Information. Ther. Innov. Regul. Sci. 59, 288–294 (2025).

64. Nelson, S. J., Zeng, K., Kilbourne, J., Powell, T. & Moore, R. Normalized names for clinical drugs: RxNorm at 6 years. J. Am. Med. Informatics Assoc. 18, 441–448 (2011).

65. WHO Collaborating Centre for Drug Statistics Methodology. Guidelines for ATC Classification and DDD Assignment. (Oslo, Norway, 2023).

66. Wang, C. S. et al. Detecting potential adverse drug reactions using a deep neural network model. J. Med. Internet Res. 21, (2019).

67. IMI PROTECT ADR database developed by EMA. https://web.archive.org/web/20220306092012/ http://www.imi-protect.eu/adverseDrugReactions.shtml.

68. IMI PROTECT ADR database developed by EMA. http://imi-protect.eu/adverseDrugReactions.shtml http://www.imiprotect.eu/adverseDrugReactions.shtml (2017).

69. National Institute for Health and Care Excellence. BNF: British National Formulary. 2018 https://bnf.nice.org.uk/.

70. Le-Khac, P. H., Healy, G. & Smeaton, A. F. Contrastive Representation Learning: A Framework and Review. IEEE Access 8, 193907–193934 (2020).

71. Musgrave, K., Belongie, S. & Lim, S. N. A Metric Learning Reality Check. In Lecture Notes in Computer Science (including subseries Lecture Notes in Artificial Intelligence and Lecture Notes in Bioinformatics) vol. 12370 LNCS 681–699 (Springer Science and Business Media Deutschland GmbH, 2020).

72. Pennington, J., Socher, R. & Manning, C. D. GloVe: Global Vectors for Word Representation. in Empirical Methods in Natural Language Processing (EMNLP) 1532–1543 (2014).

73. Kalyan, K. S. & Sangeetha, S. SECNLP: A survey of embeddings in clinical natural language processing. J. Biomed. Inform. 101, 103323 (2020).

74. Sohn, K. Improved deep metric learning with multi-class N-pair loss objective. In Advances in Neural Information Processing Systems (eds. D. Lee, M. Sugiyama, U. Luxburg, I. Guyon & R. Garnett) vol. 29 1857–1865 (Curran Associates, Inc., 2016).

75. Oord, A. van den, Li, Y. & Vinyals, O. Representation Learning with Contrastive Predictive Coding. (2018).

76. Mikolov, T., Chen, K., Corrado, G. & Dean, J. Efficient estimation of word representations in vector space. in 1st International Conference on Learning Representations, ICLR 2013 - Workshop Track Proceedings (International Conference on Learning Representations, ICLR, 2013).

77. Clarke, K. R., Somerfield, P. J. & Chapman, M. G. On resemblance measures for ecological studies, including taxonomic dissimilarities and a zero-adjusted Bray-Curtis coefficient for denuded assemblages. in Journal of Experimental Marine Biology and Ecology vol. 330 55–80 (Elsevier, 2006).

78. Kumar V., Chhabra J.K. & Dinesh K. Performance evaluation of distance metrics in the clustering algorithms. Infocomp 13, 38–51 (2014).

79. Legendre, P. & Legendre, L. Numerical Ecology. in Encyclopedia of Ecology: Volume 1-4, Second Edition vol. 3 487–493 (Elsevier, 2019).

80. Linderman, G. C. & Steinerberger, S. Clustering with t-SNE, Provably. SIAM J. Math. Data Sci. 1, 313–332 (2019).

81. Pedregosa, F. et al. Scikit-learn: Machine Learning in Python. J. Mach. Learn. Res. 12, 2825–2830 (2011).

82. Benjamini, Y. & Hochberg, Y. Controlling the False Discovery Rate: A Practical and Powerful Approach to Multiple Testing. J. R. Stat. Soc. Ser. B Stat. Methodol. 57, 289–300 (1995).

83. Chollet, F. & others. Keras. at (2015).

84. Inc., P. T. Collaborative data science. at https://plot.ly (2015).

85. Wolf, T. et al. Transformers: State-of-the-Art Natural Language Processing. EMNLP 2020 - Conf. Empir. Methods Nat. Lang. Process. Proc. Syst. Demonstr. 38–45 (2020) doi:10.18653/V1/2020.EMNLP-DEMOS.6.

86. Paszke, A. et al. PyTorch: An Imperative Style, High-Performance Deep Learning Library. Adv. Neural Inf. Process. Syst. 32, (2019).

