## Supplementary information for "Contrastive learning of adverse events to provide effective and interpretable vector representations for machine-assisted pharmacovigilance"

### 1. Visualizations of embeddings

#### 1.1. NSG-reaction2reaction

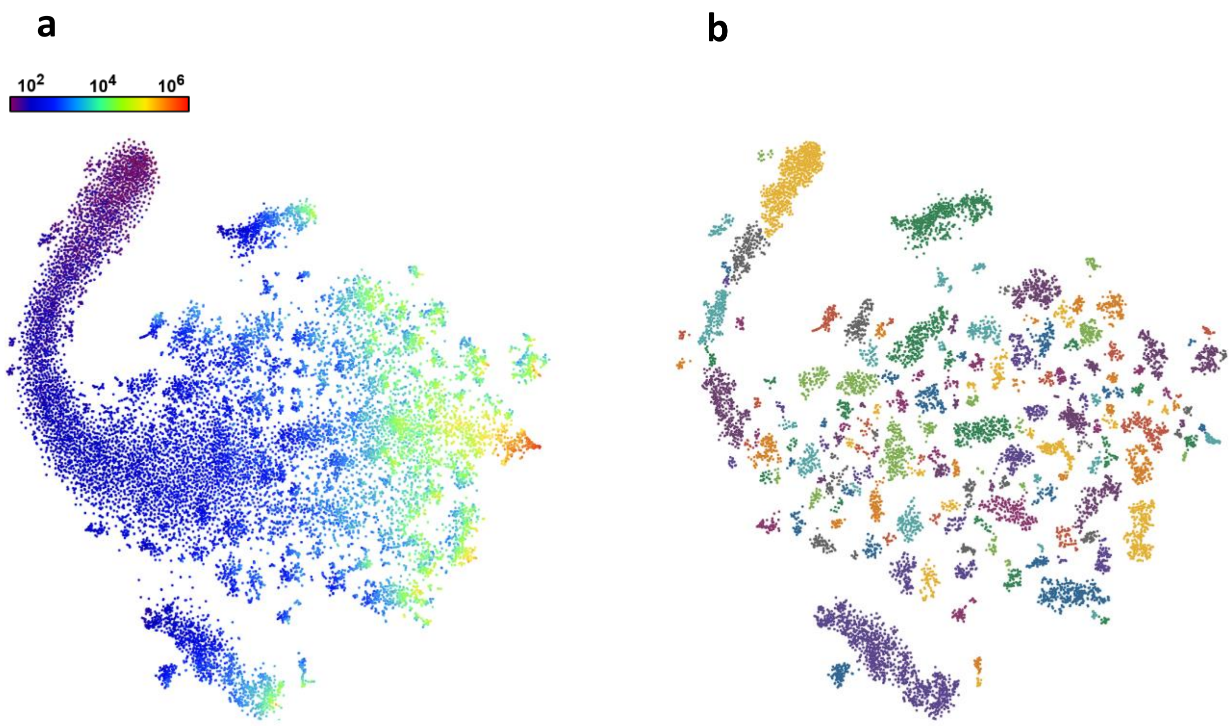

***Supplementary Figure 1.*** *t-SNE plot made from the embedding results of the NSG model with reaction2reaction context-sampling applying logarithmic frequency-based coloring* ***(a)*** *and HDBSCAN clustering-based coloring, with noise removed* ***(b)***

### 1.2. NSG-reaction2all

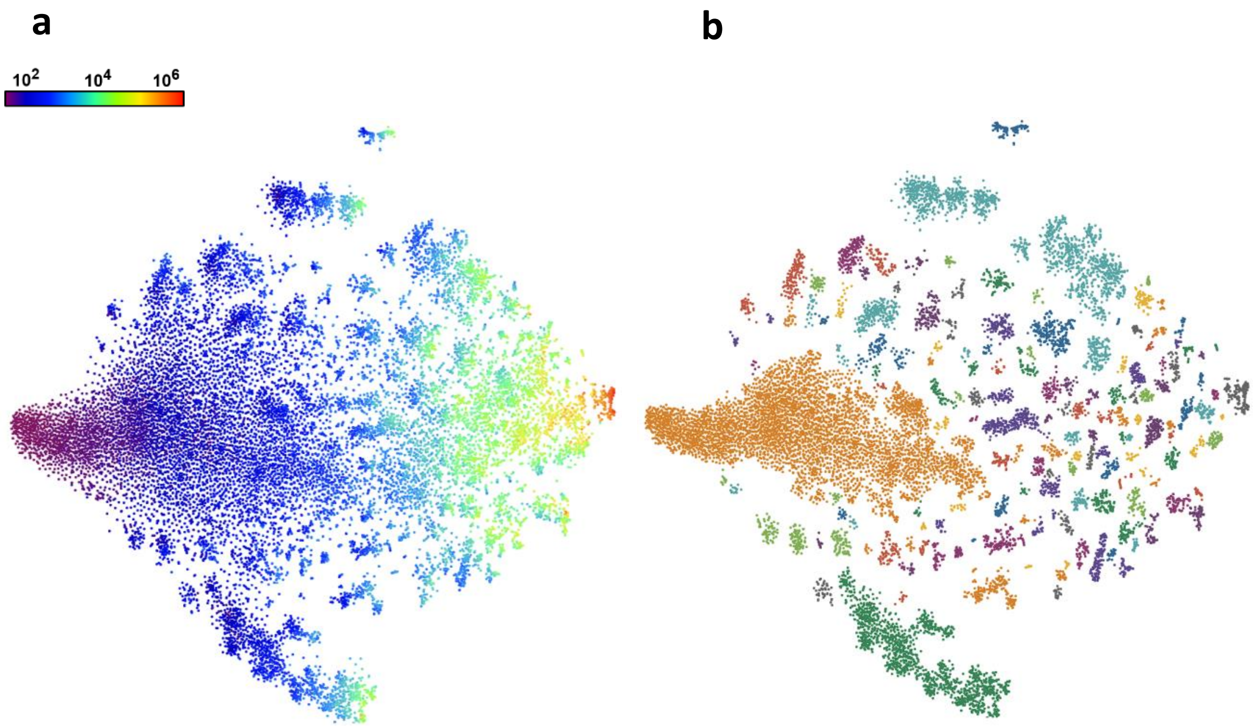

***Supplementary Figure 2.*** *t-SNE plot made from the embedding results of the NSG model with reaction2all context-sampling applying logarithmic frequency-based coloring* ***(a)*** *and HDBSCAN clustering-based coloring, with noise removed* ***(b)***

### 1.3. NTX-reaction2drug

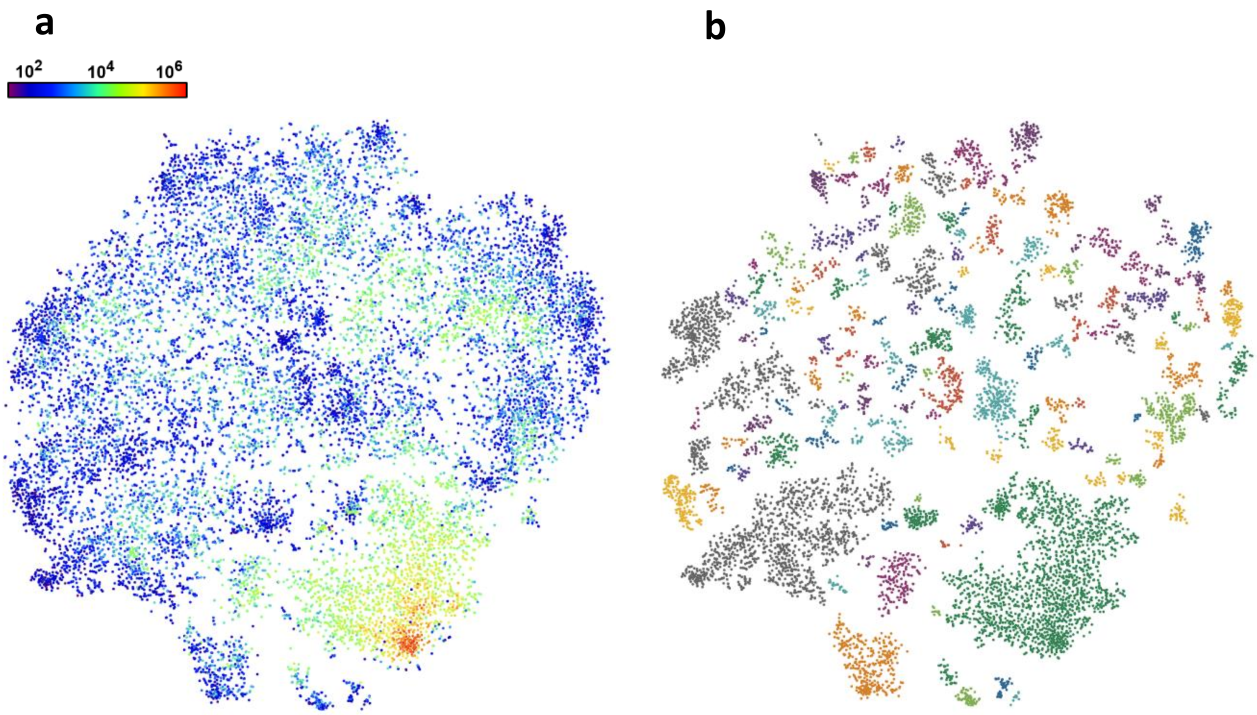

***Supplementary Figure 3.*** *t-SNE plot made from the embedding results of the NTX model with reaction2drug context-sampling applying logarithmic frequency-based coloring* ***(a)*** *and HDBSCAN clustering-based coloring, with noise removed* ***(b)***

### 1.4. NTX-reaction2reaction

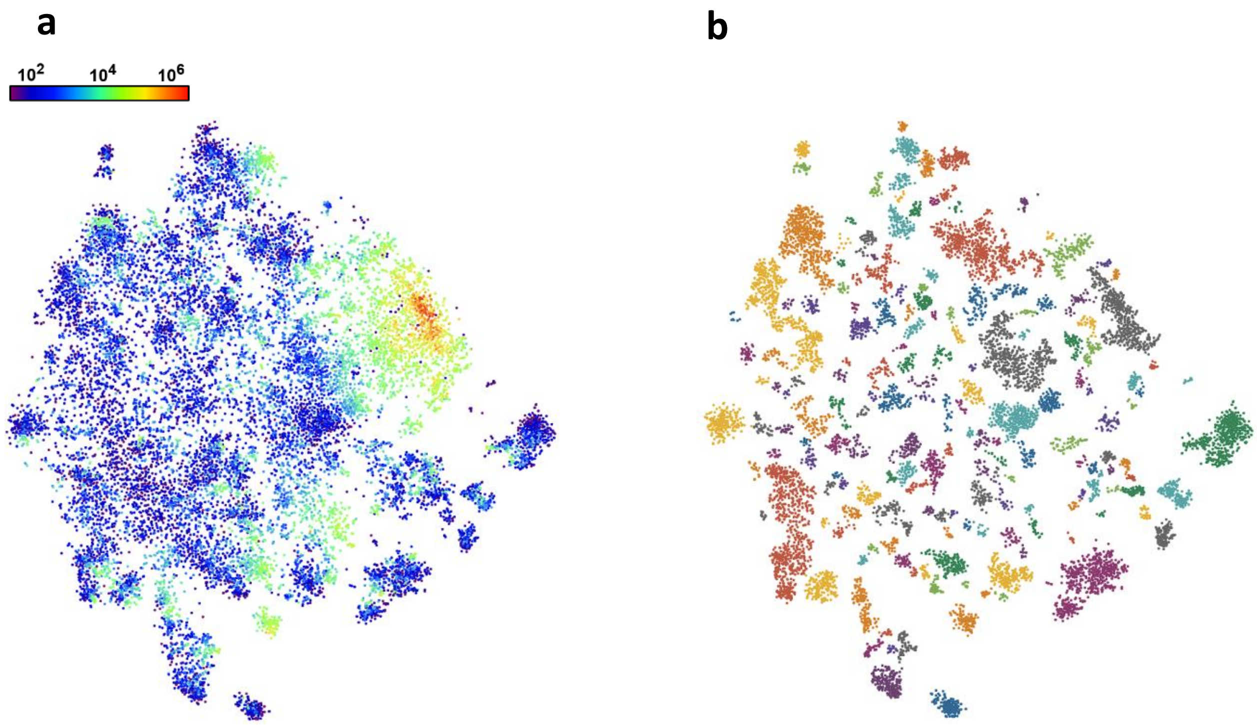

***Supplementary Figure 4.*** *t-SNE plot made from the embedding results of the NTX model with reaction2reaction context-sampling applying logarithmic frequency-based coloring* ***(a)*** *and HDBSCAN clustering-based coloring, with noise removed* ***(b)***

### 1.5. NTX-reaction2all

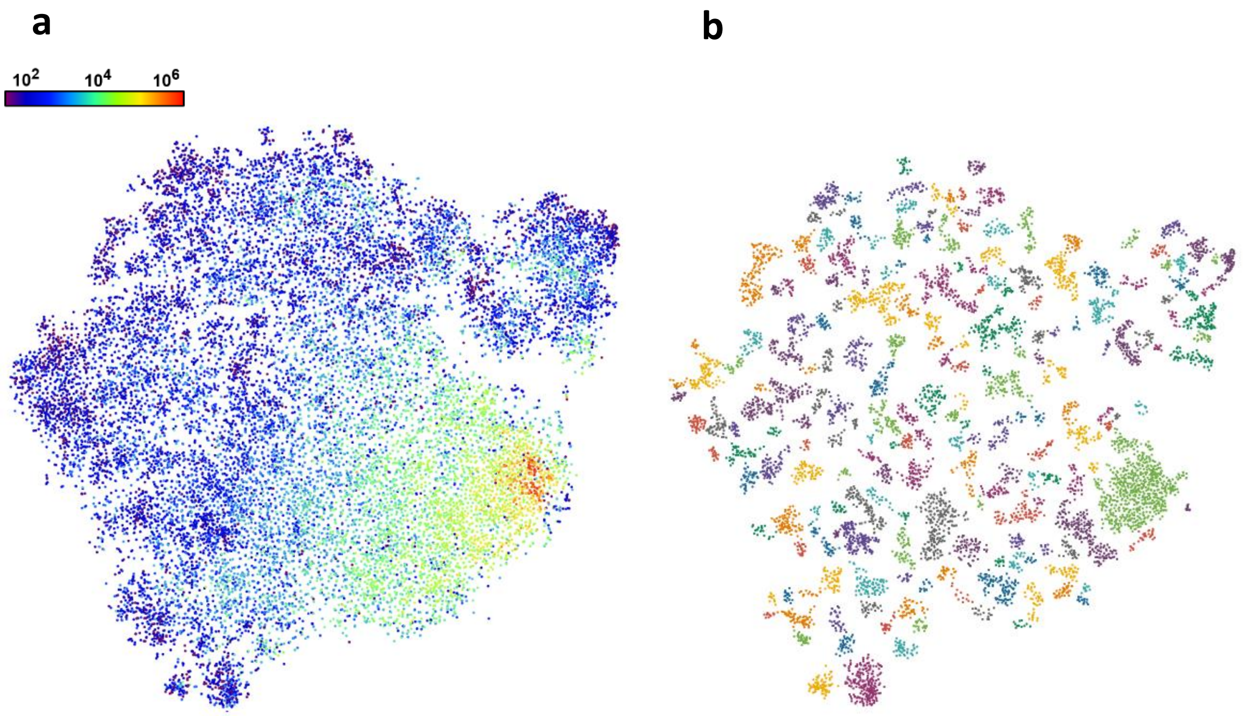

***Supplementary Figure 5.*** *t-SNE plot made from the embedding results of the NTX model with reaction2all context-sampling applying logarithmic frequency-based coloring* ***(a)*** *and HDBSCAN clustering-based coloring, with noise removed* ***(b)***

### 2. Top 10 closest neighbors

#### 2.1. NSG-reaction2drug

***Supplementary Table 1.*** *Top 10 closest neighbors of the MedDRA preferred term Myocardial Ischeamia based on various similarity and distance measures in the embedding space learnt by the NSG model with reaction2drug context-sampling*

| | **Cosine similarity** | **Euclidean distance** | **Dot product** | **Bray-Curtis dissimilarity** | **Canberra distance** |
|---|---|---|---|---|---|
| 1 | Acute Coronary Syndrome | Acute Coronary Syndrome | End Stage Renal Disease | Acute Coronary Syndrome | Acute Coronary Syndrome |
| 2 | Angina Pectoris | Electrocardiogram Abnormal | Bone Density Decreased | Electrocardiogram Abnormal | Coronary Artery Disease |
| 3 | Coronary Artery Disease | Hypotension | Device Expulsion | Hypotension | Cardiac Failure |
| 4 | Acute Myocardial Infarction | Cardiomegaly | Drug Withdrawal Syndrome Neonatal | Mitral Valve Incompetence | Electrocardiogram Abnormal |
| 5 | Arrhythmia | Supraventricular Tachycardia | Renal Injury | Arteriosclerosis | Hypokinesia |
| 6 | Circulatory Collapse | Arteriosclerosis | Colorectal Cancer | Hypokinesia | Angina Pectoris |
| 7 | Cardiac Failure | Chest Pain | Learning Disability | Chest Pain | Chest Pain |
| 8 | Electrocardiogram Abnormal | Ventricular Extrasystoles | Renal Cancer | Cardiomegaly | Fall |
| 9 | Shock | Malnutrition | Bladder Cancer | Circulatory Collapse | Pericardial Effusion |
| 10 | Mitral Valve Incompetence | Fall | Bone Loss | Angina Pectoris | Arrhythmia |

### 2.2. NSG-reaction2reaction

***Supplementary Table 2.*** *Top 10 closest neighbors of the MedDRA preferred term Myocardial Ischeamia based on various similarity and distance measures in the embedding space learnt by the NSG model with reaction2reaction context-sampling*

| | **Cosine similarity** | **Euclidean distance** | **Dot product** | **Bray-Curtis dissimilarity** | **Canberra distance** |
|---|---|---|---|---|---|
| 1 | Left Ventricular Dysfunction | Ischaemia | Skeletal Injury | Left Ventricular Dysfunction | Ischaemia |
| 2 | Ejection Fraction Decreased | Left Ventricular Dysfunction | Device Deployment Issue | Ischaemia | Peripheral Vascular Disorder |
| 3 | Ischaemia | Ejection Fraction Decreased | Colorectal Cancer | Ejection Fraction Decreased | Ejection Fraction Decreased |
| 4 | Acute Myocardial Infarction | Acute Myocardial Infarction | Multiple Fractures | Acute Myocardial Infarction | Sinus Bradycardia |
| 5 | Ventricular Hypokinesia | Ventricular Extrasystoles | Foreign Body In Reproductive Tract | Supraventricular Tachycardia | Pharyngitis |
| 6 | Ventricular Extrasystoles | Supraventricular Tachycardia | Complication Of Device Removal | Ventricular Extrasystoles | Troponin Increased |
| 7 | Sinus Bradycardia | Cardiomyopathy | Osteonecrosis | Generalised Oedema | Stress Cardiomyopathy |
| 8 | Supraventricular Tachycardia | Cardiac Valve Disease | Device Expulsion | Ventricular Hypokinesia | Ventricular Tachycardia |
| 9 | Cardiomyopathy | Peripheral Vascular Disorder | Application Site Erythema | Peripheral Vascular Disorder | Acute Myocardial Infarction |
| 10 | Peripheral Vascular Disorder | Generalised Oedema | Sprue-Like Enteropathy | Ventricular Tachycardia | Electrocardiogram St Segment Depression |

### 2.3. NSG-reaction2all

***Supplementary Table 3.*** *Top 10 closest neighbors of the MedDRA preferred term Myocardial Ischeamia based on various similarity and distance measures in the embedding space learnt by the NSG model with reaction2all context-sampling*

| | **Cosine similarity** | **Euclidean distance** | **Dot product** | **Bray-Curtis dissimilarity** | **Canberra distance** |
|---|---|---|---|---|---|
| 1 | Arteriosclerosis Coronary Artery | Acute Coronary Syndrome | Colorectal Cancer | Left Ventricular Dysfunction | Left Ventricular Dysfunction |
| 2 | Sinus Tachycardia | Ejection Fraction Decreased | Oesophageal Carcinoma | Arteriosclerosis Coronary Artery | Atrioventricular Block |
| 3 | Left Ventricular Dysfunction | Ventricular Arrhythmia | Bladder Cancer | Atrioventricular Block | Sinus Tachycardia |
| 4 | Ventricular Extrasystoles | Cholelithiasis | Dysphoria | Pericardial Effusion | Ventricular Extrasystoles |
| 5 | Acute Coronary Syndrome | Sinus Tachycardia | Gastric Cancer | Sinus Bradycardia | Pericardial Effusion |
| 6 | Cholelithiasis | Troponin Increased | Hepatic Cancer | Ejection Fraction Decreased | Arteriosclerosis Coronary Artery |
| 7 | Mitral Valve Incompetence | Left Ventricular Dysfunction | Prostate Cancer | Ventricular Arrhythmia | Circulatory Collapse |
| 8 | Ejection Fraction Decreased | Atrioventricular Block | Affect Lability | Ventricular Extrasystoles | Sinus Bradycardia |
| 9 | Bundle Branch Block Right | Sinus Bradycardia | Renal Cancer | Cholelithiasis | Atrioventricular Block First Degree |
| 10 | Sinus Bradycardia | Arteriosclerosis Coronary Artery | Psychological Trauma | Sinus Tachycardia | Ecchymosis |

### 2.4. NTX-reaction2drug

***Supplementary Table 4.*** *Top 10 closest neighbors of the MedDRA preferred term Myocardial Ischeamia based on various similarity and distance measures in the embedding space learnt by the NTX model with reaction2drug context-sampling*

| | **Cosine similarity** | **Euclidean distance** | **Dot product** | **Bray-Curtis dissimilarity** | **Canberra distance** |
|---|---|---|---|---|---|
| 1 | Diabetic Ketoacidosis | Diabetic Ketoacidosis | Sitophobia | Diabetic Ketoacidosis | Rales |
| 2 | Subdural Haematoma | Subdural Haematoma | Hypoxia | Subdural Haematoma | Pneumothorax |
| 3 | Aspiration | Diverticulum | White Blood Cell Count Increased | Rales | Subdural Haematoma |
| 4 | Blood Uric Acid Increased | Large Intestine Polyp | Electrocardiogram Qt Prolonged | Aspiration | Ileus |
| 5 | Pancytopenia | Sudden Death | Prone Position | Pancytopenia | Diabetic Ketoacidosis |
| 6 | Sudden Death | Infarction | Pulmonary Hypertension | Sudden Death | Clostridium Difficile Infection |
| 7 | International Normalised Ratio Increased | Renal Injury | Renal Impairment | Pneumothorax | Acute Respiratory Distress Syndrome |
| 8 | Haemodialysis | Ischaemia | Discomfort | Blood Uric Acid Increased | Polyneuropathy |
| 9 | Haematocrit Decreased | Monoplegia | Myocardial Infarction | Haemodialysis | Pancytopenia |
| 10 | Diverticulum | Blood Phosphorus Decreased | Aspiration | Diverticulum | Blood Phosphorus Decreased |

### 2.5. NTX-reaction2reaction

***Supplementary Table 5.*** *Top 10 closest neighbors of the MedDRA preferred term Myocardial Ischeamia based on various similarity and distance measures in the embedding space learnt by the NTX model with reaction2reaction context-sampling*

| | **Cosine similarity** | **Euclidean distance** | **Dot product** | **Bray-Curtis dissimilarity** | **Canberra distance** |
|---|---|---|---|---|---|
| 1 | Sinus Tachycardia | Atrioventricular Block First Degree | Mitral Valve Incompetence | Sinus Tachycardia | Sinus Tachycardia |
| 2 | Bundle Branch Block Right | Sinus Tachycardia | Foot Deformity | Coronary Artery Disease | Coronary Artery Disease |
| 3 | Mitral Valve Incompetence | Left Ventricular Hypertrophy | Sacral Hypoplasia | Sinus Bradycardia | Pulmonary Hypertension |
| 4 | Left Ventricular Hypertrophy | Sinus Bradycardia | Fallopian Tube Cancer Stage Iv | Hyperkalaemia | Arteriosclerosis Coronary Artery |
| 5 | Hyperkalaemia | Hypovolaemia | Bundle Branch Block Right | Left Ventricular Hypertrophy | Hyperlipidaemia |
| 6 | Sinus Bradycardia | Atrial Flutter | Tracheal Fistula Repair | Bundle Branch Block Right | Respiratory Disorder |
| 7 | Coronary Artery Disease | Ventricular Tachycardia | Macula Thickness Measurement | Arteriosclerosis Coronary Artery | Hyperkalaemia |
| 8 | Bundle Branch Block Left | Arterial Occlusive Disease | Fall | Bundle Branch Block Left | Femur Fracture |
| 9 | Hypovolaemia | Hyperkalaemia | Oesophageal Carcinoma Stage 0 | Hypovolaemia | Sinus Bradycardia |
| 10 | Arteriosclerosis Coronary Artery | Bundle Branch Block Right | Tricuspid Valve Incompetence | Atrioventricular Block First Degree | Hyperglycaemia |

### 2.6. NTX-reaction2all

***Supplementary Table 6.*** *Top 10 closest neighbors of the MedDRA preferred term Myocardial Ischeamia based on various similarity and distance measures in the embedding space learnt by the NTX model with reaction2all context-sampling*

| | **Cosine similarity** | **Euclidean distance** | **Dot product** | **Bray-Curtis dissimilarity** | **Canberra distance** |
|---|---|---|---|---|---|
| 1 | Increased Appetite | Increased Appetite | Hypotension | Increased Appetite | Plantar Fasciitis |
| 2 | Injection Site Bruising | Injection Site Bruising | Biopsy Rectum Abnormal | Plantar Fasciitis | Liver Disorder |
| 3 | Polydipsia | Neck Injury | Intervertebral Disc Degeneration | Injection Site Bruising | Bone Lesion |
| 4 | Mood Swings | Plantar Fasciitis | Femur Fracture | Muscle Disorder | Hallucination, Visual |
| 5 | Barrett's Oesophagus | Barrett's Oesophagus | Malaise | Joint Range Of Motion Decreased | Neuritis |
| 6 | Neck Injury | Mood Swings | Hypertension | Liver Disorder | Increased Appetite |
| 7 | Wound Complication | Skin Necrosis | Condition Aggravated | Neuritis | Joint Range Of Motion Decreased |
| 8 | Joint Range Of Motion Decreased | Wound Complication | Neuropathy Peripheral | Barrett's Oesophagus | Wound Complication |
| 9 | Plantar Fasciitis | Chronic Sinusitis | Rash | Wound Complication | Blood Calcium Decreased |
| 10 | Mental Status Changes | Tenderness | Discomfort | Mental Status Changes | Muscle Disorder |

### 2.7. MedDRA

***Supplementary Table 7.*** *Top 10 closest neighbors of the MedDRA preferred term Myocardial Ischeamia based on various node-based hierarchical similarity measures*[1] *calculated in the MedDRA terminology network*

| | **Mean minimum path** | **Resnik similarity** | **Jiang-Conrath similarity** | **Lin similarity** |
|---|---|---|---|---|
| 1 | Acute Coronary Syndrome | Acute Coronary Syndrome | Myocardial Infarction | Myocardial Infarction |
| 2 | Acute Myocardial Infarction | Acute Myocardial Infarction | Coronary Artery Disease | Coronary Artery Disease |
| 3 | Angina Pectoris | Angina Pectoris | Angina Pectoris | Angina Pectoris |
| 4 | Angina Unstable | Angina Unstable | Acute Myocardial Infarction | Acute Myocardial Infarction |
| 5 | Anginal Equivalent | Anginal Equivalent | Arteriosclerosis Coronary Artery | Arteriosclerosis Coronary Artery |
| 6 | Cardiac Perfusion Defect | Cardiac Perfusion Defect | Acute Coronary Syndrome | Acute Coronary Syndrome |
| 7 | Chronic Coronary Syndrome | Chronic Coronary Syndrome | Angina Unstable | Angina Unstable |
| 8 | Coronary No-Reflow Phenomenon | Coronary No-Reflow Phenomenon | Arteriosclerosis | Stress Cardiomyopathy |
| 9 | Coronary Steal Syndrome | Coronary Steal Syndrome | Aortic Arteriosclerosis | Ischaemic Cardiomyopathy |
| 10 | Microvascular Coronary Artery Disease | Microvascular Coronary Artery Disease | Stress Cardiomyopathy | Prinzmetal Angina |

### 3. Top 10 closest neighbors categorization method

***Supplementary Table 8.*** *The exact sections of the international guidelines we utilized as reference for the evaluation of the top 10 neighbors of the MedDRA preferred term Myocardial Ischeamia, and the resulting categories. Myocardial Ischaemia has a major role in two syndromes, namely Acute Coronary Syndrome (ACS) and Chronic Coronary Syndrome (CCS), which were used as starting points for many of the evaluations.*

| Reaction | Category | Reference |
|---|---|---|
| Acute Coronary Syndrome | **Possible complication** | • ESC Guidelines for ACS[2]: 2.1<br>• ESC Guidelines for CCS[3]: 2 |
| Acute Myocardial Infarction | **Possible complication** | • ESC Guidelines for ACS[2]: 2.1 as ñmyocardial infarctionò<br>• ESC Guidelines for CCS[3], supp.: 1.1.1.2 |
| Angina Pectoris | **Symptom** | • ESC Guidelines for ACS[2]: 2.1<br>• ESC Guidelines for CCS[3]: 2.2 |
| Arrhythmia | **Possible complication** | • ESC Guidelines for ACS[2]: 12.2.5<br>• ESC Guidelines for CCS[3]: 3.3.5, 6.3 |
| Arteriosclerosis Coronary Artery | **Risk factor** | • ESC Guidelines for CCS[3]: 2.1<br>• ESC Guidelines for ACS[2]: 12.1 as ñatherosclerosisò<br>• Definition of myocardial infarction[4]: 7.1, 7.2 as ñatherosclerotic plaqueò |
| Aspiration | **Indirect connection** | *Aspiration* is a potential consequence of any surgery, including heart surgery due to *Myocardial Ischaemia*. *Aspiration* is also a potential consequence of loss of consciousness from *Circulatory Collapse*. |
| Barettôs Oesophagus | **No connection** | *Angina Pectoris* (main symptom of *Myocardial Ischaemia*) is one of the symptoms of *Barett's Oesophagus*. |
| Blood Uric Acid Increased | **Shared pathophysiological cause** (chronic kidney disease) | • Clinical Practice Guideline for the Evaluation and Management of Chronic Kidney Disease[5]: 3.14 Hyperuricemia<br>• ESC Guidelines for CCS[3]: 5.3.8 |
| Bundle Branch Block Left | **Sign** | • ESC Guidelines for ACS[2]: 3.2.1 as ñright bundle branch blockò<br>• ESC Guidelines for ACS[2], supp.: 3.2.1 as ñright bundle branch blockò<br>• Definition of myocardial infarction[4]: 32 |
| Bundle Branch Block Right | **Sign** | |
| Cardiac Failure | **Possible complication** | • ESC Guidelines for CCS[3]: 3.3.5, 6.3 as ñheart failureò<br>• ESC Guidelines for CCS[3], supp.: 1.1.1.3 as ñheart failureò<br>• ESC Guidelines for ACS[2]: 12.2.1 as ñheart failureò |
| Cardiomyopathy | **Possible complication, Risk factor** | Complication:<br>• ESC Guidelines for CCS[3]: 4.4.2 as ñischaemic cardiomyopathyò<br>Risk factor:<br>• see Left Ventricular Hypertrophic Cardiomyopathy |
| Coronary Artery Disease | **Risk factor** | • ESC Guidelines for CCS[3]: 2<br>• Definition of myocardial infarction[4]: 7.1 |
| Cholelithiasis | **Shared pathophysiological cause** (obesity, diabetes mellitus, dyslipidemia, sedentary lifestyle) | • ASL Clinical Practice Guidelines on the prevention, diagnosis and treatment of gallstones: Prevention of gallstones: Primary prevention of gallstones[6]<br>• obesity ï ESC Guidelines for CCS[3]: 4.1.2.2<br>• diabetes mellitus ï ESC Guidelines for CCS[3]: 3.3.5<br>• dyslipidemia ï ESC Guidelines for CCS[3]: 4.3.2<br>• sedentary lifestyle ï ESC Guidelines for CCS[3]: 4.1.2.5 |

| | | |
|---|---|---|
| Circulatory Collapse | **Possible complication** | • ESC Guidelines for ACS[2]: 3.1.2 as ñcirculatory compromiseò<br>• ESC Guidelines for ACS[2]: 12.2.6 as ñcardiocirculatory collapseò |
| Diabetic Ketoacidosis | **Shared pathophysiological cause** (diabetes mellitus) | • Standards of Care in Diabetesð 2025: 6. Glycemic Goals and Hypoglycemia: HYPERGLYCEMIC CRISES: DIAGNOSIS, MANAGEMENT, AND PREVENTION (or supplementary page: S139)[7]<br>• ESC Guidelines for CCS[3]: 3.3.5 |
| Diverticulum | **No connection** | *Diverticulum* might be present due to its frequent co-occurrences with *Myocardial Ischaemia* in the elderly. |
| Ejection Fraction Decreased | **Sign** | • ESC Guidelines for CCS[3]: 3.2.2<br>• ESC Guidelines for CCS[3], supp.: 1.1.1.3 as ñleft ventricular ejection fractionò |
| Electrocardiogram Abnormal | **Sign** | • ESC Guidelines for ACS[2]: 2.1 as ñECG changesò<br>• ESC Guidelines for CCS[3]: 3.1.2.1 |
| Haematocrit Decreased | **Shared pathophysiological cause** (chronic kidney disease) | • KDIGO Clinical Practice Guideline[8]: Chapter 1: Testing for Anemia: Rationale<br>• ESC Guidelines for CCS[3]: 5.3.8 |
| Haemodialysis | **Shared pathophysiological cause** (chronic kidney disease) | • Clinical Practice Guideline for the Evaluation and Management of Chronic Kidney Disease[5]: Heamodialysis a general kidney replacement therapy in chronic kidney disease<br>• ESC Guidelines for CCS[3]: 5.3.8 |
| Hyperkalaemia | **Shared pathophysiological cause** (chronic kidney disease) | • Clinical Practice Guideline for the Evaluation and Management of Chronic Kidney Disease[5]: Chapter 2: Practice Point 2.2.2<br>• ESC Guidelines for CCS[3]: 5.3.8 |
| Hypovolaemia | **Shared pathophysiological cause** (diabetes mellitus) | • Standards of Care in Diabetesð 2025: 6. Glycemic Goals and Hypoglycemia[7]: HYPERGLYCEMIC CRISES: DIAGNOSIS, MANAGEMENT, AND PREVENTION (or supplementary page: S140)[7]<br>• ESC Guidelines for CCS[3]: 3.3.5 |
| INR Increased | **No connection** | Increased INR (international normalised ratio) is not strictly related:<br>• ESC Guidelines for CCS[3]: 4.3.1.2<br>And common:<br>• ESC Guidelines for ACS[2]: 6.5.1 |
| Increased Appetite | **Shared pathophysiological cause** (diabetes mellitus) | • Standards of Care in Diabetesð 2025: 6. Glycemic Goals and Hypoglycemia[7]: HYPOGLYCEMIA ASSESSMENT, PREVENTION, AND TREATMENT: Hypoglycemia Definitions and Event Rates<br>• ESC Guidelines for CCS[3]: 3.3.5 |
| Injection Site Bruising | **No connection** | Generally, due to high INR, which is frequent and not specific to *Myocardial Ischaemia*. |
| Ischaemia | **Synonym** | Less specific term, commonly used referring to *Myocardial Ischaemia* in a cardiological context. |
| Joint Range of Motion Decreased | **No connection** | *Joint Range of Motion Decreased* might be present due to its frequent co-occurrences with *Myocardial Ischaemia* in the elderly. |
| Left Ventricular Dysfunction | **Sign** | • ESC Guidelines for CCS[3]: 3.2.2 as ñdecreased LV functionò |
| Left Ventricular Hypertrophy | **Risk factor** | • ESC Guidelines for CCS[3]: 2.1 as ñmyocardial hypertrophyò<br>• Definition of myocardial infarction[4]: 34.1 ñhypertrophic cardiomyopathyò |
| Mental Status Changes | **Possible complication, Risk factor** | Depression:<br>• ESC Guidelines for CCS[3]: 4.1.2.4<br>• ESC Guidelines for ACS[2]: 3.2.4 |

| | | |
|---|---|---|
| Mitral Valve Incompetence | **Possible complication** | • ESC Guidelines for CCS[3]: 3.3.5, 6.3<br>• ESC Guidelines for CCS[3], supp.: 1.1.1.4 as ñmitral valve regurgitationò<br>• ESC Guidelines for ACS[2]: 12.2.2 as ñacute mitral regurgitationò |
| Mood Swings | **No connection** | No information supporting any connection was found. |
| Neck Injury | **No connection** | Potential consequence of falling, caused by *Circulatory Collapse*. |
| Pancytopenia | **No connection** | No information supporting any connection was found. |
| Peripheral Vascular Disorder | **Shared pathophysiological cause** (diabetes mellitus) | • Standards of Care in Diabetesð 2025: 12. Retinopathy, Neuropathy, and Foot Care (or supplementary page S259)[9]<br>• ESC Guidelines for CCS[3]: 3.3.5 |
| Plantar Fasciitis | **No connection** | No information supporting any connection was found. |
| Polydipsia | **Shared pathophysiological cause** (diabetes mellitus) | • Standards of Care in Diabetesð 2025: 2. Diagnosis and Classification of Diabetes: Diagnostic Tests for Diabetes[10]<br>• ESC Guidelines for CCS[3]: 3.3.5 |
| Shock | **Possible complication** | • ESC Guidelines for ACS[2]: 2.1 as ñcardiac complicationò |
| Sinus Bradycardia | **Possible complication, Risk factor** | Complication:<br>• ESC Guidelines for ACS[2]: Recommendation Table 14 as ñbradyarrhthmiasòor ñsinus bradycardiaò<br>Risk factor:<br>• ESC Guidelines for ACS[2]: 12.1 at ñnon-coronary mechanismsò |
| Sinus Tachycardia | **Risk factor** | • ESC Guidelines for CCS[3]: 2.1 as ñtachycardiaò<br>• Definition of myocardial infarction[4]: 33 |
| Subdural Haematoma | **Indirect connection** | *Subdural Haematoma* is bleeding in the skull, typically due to head trauma, which might be caused by fainting and/or falling after a *Circulatory Collapse*. |
| Sudden Death | **Possible complication** | • ESC Guidelines for CCS[3]: 3.3.5 as ñcardiovascular deathò |
| Supraventricular Tachycardia | **Possible complication** | • ESC Guidelines for ACS[2]: 12.2.5.1 as a subtype, ñatrial fibrillationò |
| Ventricular Extrasystoles | **Possible complication** | • ESC Guidelines for CCS[3], supp.: 1.1.1.3 as ñventricular arrhythmiaòwhich is a broader term including *Ventricular Extrasystoles*<br>• ESC Guidelines for ACS[2]: 12.2.5.2 as ñventricular premature beatsò |
| Ventricular Hypokinesia | **Sign** | • ESC Guidelines for CCS[3]: 3.2.2 as ñregional wall motion abnormalitiesò |
| Wound Complication | **Shared pathophysiological Cause** (diabetes mellitus) | • Standards of Care in Diabetesð 2025: 13. Older Adults: Older Adults With Complications and Reduced Functionality (or supplementary page S270)[11]<br>• ESC Guidelines for CCS[3]: 3.3.5 |

148

### 4. Gain norm vs dispersion test results

***Supplementary Table 9.*** *Gain norm vs. the ratio of drugs with significantly ($q < 0.05$) different reaction embedding dispersions compared to random noise (sign. ratio), throughout our six embeddings and the four BERT-based ones, as calculated using various similarity and distance measures*

| | | Cosine similarity | Euclidean distance | Dot product | Bray-Curtis dissimilarity | Canberra distance |
|---|---|---|---|---|---|---|
| **NSG-reaction2drug** | sign. ratio | 0.9944 | 0.1098 | 0.9919 | 0.9938 | 0.9913 |
| | gain norm | 0.4482 | -0.0083 | 0.2779 | 0.3871 | 0.4620 |
| **NSG-reaction2reaction** | sign. ratio | 0.9882 | 0.0000 | 0.9919 | 0.9882 | 0.9845 |
| | gain norm | 0.3105 | -0.0004 | 0.3482 | 0.2661 | 0.3117 |
| **NSG-reaction2all** | sign. ratio | 0.9746 | 0.0496 | 0.9926 | 0.9609 | 0.9398 |
| | gain norm | 0.2544 | 0.0219 | 0.2795 | 0.2665 | 0.2139 |
| **NTX-reaction2drug** | sign. ratio | 0.9901 | 0.9895 | 0.9882 | 0.9895 | 0.9864 |
| | gain norm | 0.3150 | 0.2652 | 0.1912 | 0.1822 | 0.2868 |
| **NTX-reaction2reaction** | sign. ratio | 0.9876 | 0.9907 | 0.9851 | 0.9882 | 0.9870 |
| | gain norm | 0.2937 | 0.2308 | 0.1393 | 0.1960 | 0.2430 |
| **NTX-reaction2all** | sign. ratio | 0.9882 | 0.9541 | 0.9882 | 0.9895 | 0.9857 |
| | gain norm | 0.3318 | 0.1315 | 0.1847 | 0.1436 | 0.3295 |

| | | | | | | |
|---|---|---|---|---|---|---|
| **PubMedBERT** | sign. ratio | 0.9770 | 0.9752 | 0.9752 | 0.9783 | 0.9764 |
| | gain norm | 0.0823 | 0.0542 | 0.0800 | 0.0691 | 0.0396 |
| **BioMed-RoBERTa** | sign. ratio | 0 | 0 | 0 | 0.0887 | 0.4274 |
| | gain norm | 0.0069 | 0.0108 | -0.0078 | 0.0177 | 0.0141 |
| **Clinical ModernBERT** | sign. ratio | 0.8017 | 0.6024 | 0.7655 | 0.8449 | 0.8120 |
| | gain norm | 0.0553 | 0.0263 | 0.0709 | 0.0498 | 0.0333 |
| **BioClinical ModernBERT** | sign. ratio | 0 | 0 | 0.3127 | 0.1669 | 0.4677 |
| | gain norm | 0.0058 | 0.0133 | -0.0234 | 0.0215 | 0.0183 |

Pairwise permutation tests for homogeneity of multivariate dispersion[12,13] (modified permdisp tests) were performed between the validated and the random reaction groups of all drugs. A validated reaction group of a drug consists of reactions known to be associated with it according to validated drug–ADR (adverse drug reaction) data (see Section 4.1 of the manuscript). For each validated reaction group, a random reaction group with an equal size was sampled uniformly from the available reaction terms. As dispersion can only be measured in groups with multiple points, drugs with only a single known reaction had to be left out, resulting in 1612 groups remaining out of the original 1619. For the permutation tests, we used a maximum of 999 samples, if the total number of permutations was ≥999, otherwise we used all permutations. Due to multiple testing, permutation test *p*-values were corrected by a  the Benjamini–Hochberg procedure[14], yielding *q*-values. We summarized the results of the permutation tests for each embedding, and for each measure, as the ratio of validated reaction groups showing a significantly ($q < 0.05$) different dispersion compared to the corresponding random reaction group. For the gain norm calculation, see Section 4.8 of the manuscript.

### 5. The improved permutation procedure

(**Supplementary Note**)

As described by Gijbels and Omelka[13], the standard permutation procedure is not recommended for their proposed modified permdisp test, as it produces an exact permutation test only if the distribution of samples across the different groups is the same. To alleviate this issue, they described an improved ("centered") permutation procedure, where the Principal Coordinate Analysis (PCoA) transformed representation of the samples is centered to create a new distance matrix for the test. However, as a preliminary step, the distance matrix first has to be corrected to avoid negative eigenvalues arising with non-Euclidean measures, such as the cosine dissimilarity/distance (i.e. $1 - cosine\ similarity$). This can be achieved by the Cailliez correction method[15], which shifts the negative eigenvalues such that the largest negative eigenvalue becomes zero. Adapted from the work of Legendre and Legendre[16], we provide a streamlined step-by-step description of the Cailliez correction method, and the subsequent PCoA transformation and centering steps, reflecting our own implementation.

We start with two matrices, $A_1$ and $A_2$, given from the elements of distance matrix $D$ as:

$$a_{1_{ij}} = -0.5\, d_{ij}^2 \tag{1}$$

$$a_{2_{ij}} = -0.5\, d_{ij} \tag{2}$$

From these, we create two centered matrices, $\Delta_1$ and $\Delta_2$ by the formula:

$$\delta_{ij} = a_{ij} - \bar{a}_{i\cdot} - \bar{a}_{\cdot j} + \bar{a} \tag{3}$$

where $a_{ij}$ are the elements of $A_1$ and $A_2$ belonging to $\Delta_1$ and $\Delta_2$, respectively, with $\bar{a}_{i\cdot}$ , $\bar{a}_{\cdot j}$ and $\bar{a}$ denoting the row-wise, the column-wise and the global averages, respectively. Now, a special square matrix can be constructed:

$$M = \begin{bmatrix} 0 & 2\Delta_1 \\ -\mathrm{I} & -4\Delta_2 \end{bmatrix} \tag{4}$$

where $0$ is a null matrix and $\mathrm{I}$ is an identity matrix. Adding the largest positive eigenvalue $c$ of matrix $M$ to all non-diagonal elements of the distance matrix $D$, we modify $D$ with just the right amount, so that its eigenvalues will be non-negative:

$$\hat{d}_{hi} = d_{hi} + c \qquad \text{for } h \neq i \tag{5}$$

With this corrected distance matrix $\widehat{D}$, we begin the PCoA transformation by first creating the centered matrix $\widehat{\Delta}$ by using Eq. 3 again, now with elements from matrix $\hat{A}$, given by the transformation of $\widehat{D}$:

$$\hat{a}_{ij} = -0.5\, \hat{d}_{ij}^2 \tag{6}$$

Then, the PCoA-transformed representation $U$ of the centered matrix $\widehat{\Delta}$ is given by:

$$U = E_m \Lambda_m^{\frac{1}{2}} \quad (7)$$

where $E$ is a matrix formed by the eigenvectors of $\widehat{\Delta}$ corresponding to the $m$ largest eigenvalues in descending order (with $m$ being equal to the dimensions of $\widehat{\Delta}$ in this case) and $\Lambda$ is a diagonal matrix of the eigenvalues.

We center the representations (rows of $U$) by the geometric median of their corresponding groups (estimated by the algorithm of Vardi and Zhang[17]) via simply subtracting the median vector from each representation. A new distance matrix $\widetilde{D}$ is then created from these centered representations, using the Euclidean distance:

$$\tilde{d}_{ij} = \left\| u_i - u_j \right\|_2 \quad (8)$$

Finally, the correction value $c$ is subtracted from the non-diagonal elements of $\widetilde{D}$ to get the final adjusted distance matrix used to calculate the test statistic (see Eq. 7.1-7.3 in Section 4.7 of the manuscript).

### 6. Gain norm during training

#### 6.1. NSG-reaction2reaction

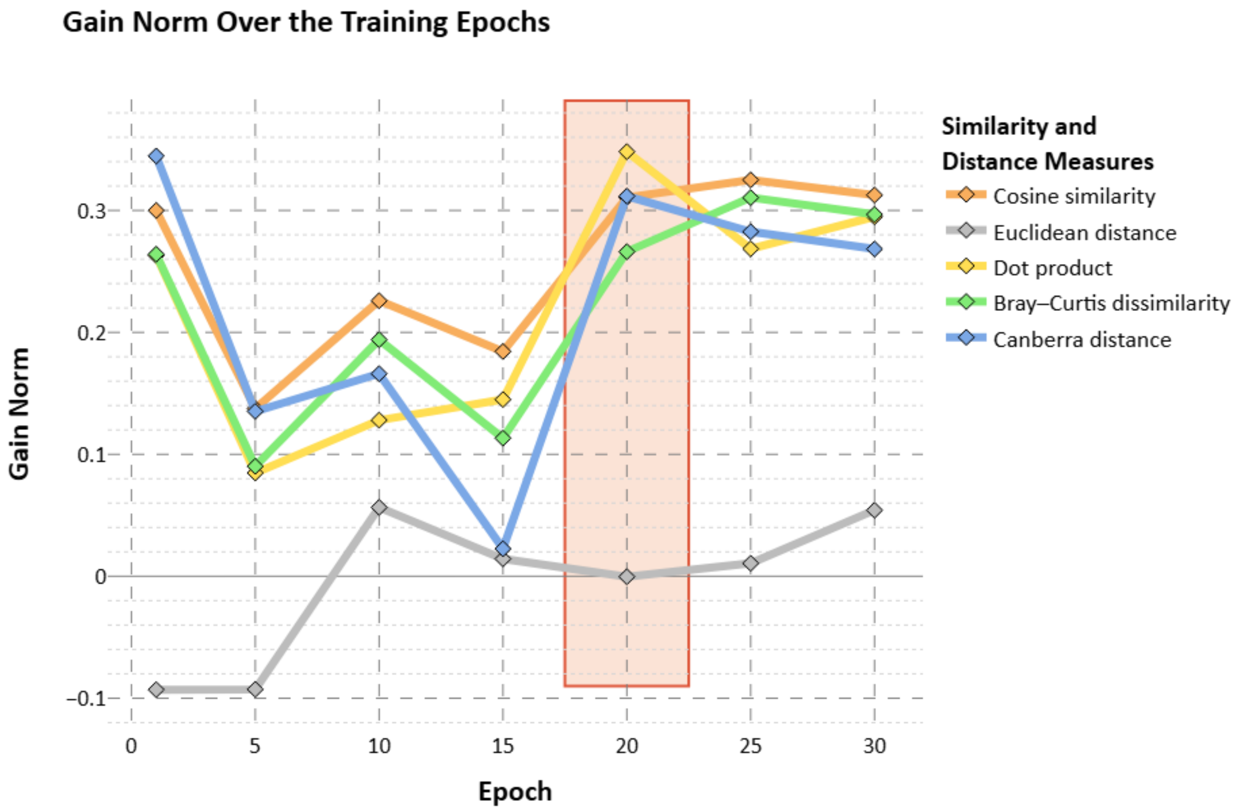

***Supplementary Figure 6.*** *Gain norm calculated with various similarity and distance measures throughout the training of the NSG model with reaction2reaction context-sampling. The epoch from which the final embedding was extracted is highlighted by a semi-transparent red rectangle.*

### 6.2. NSG-reaction2all

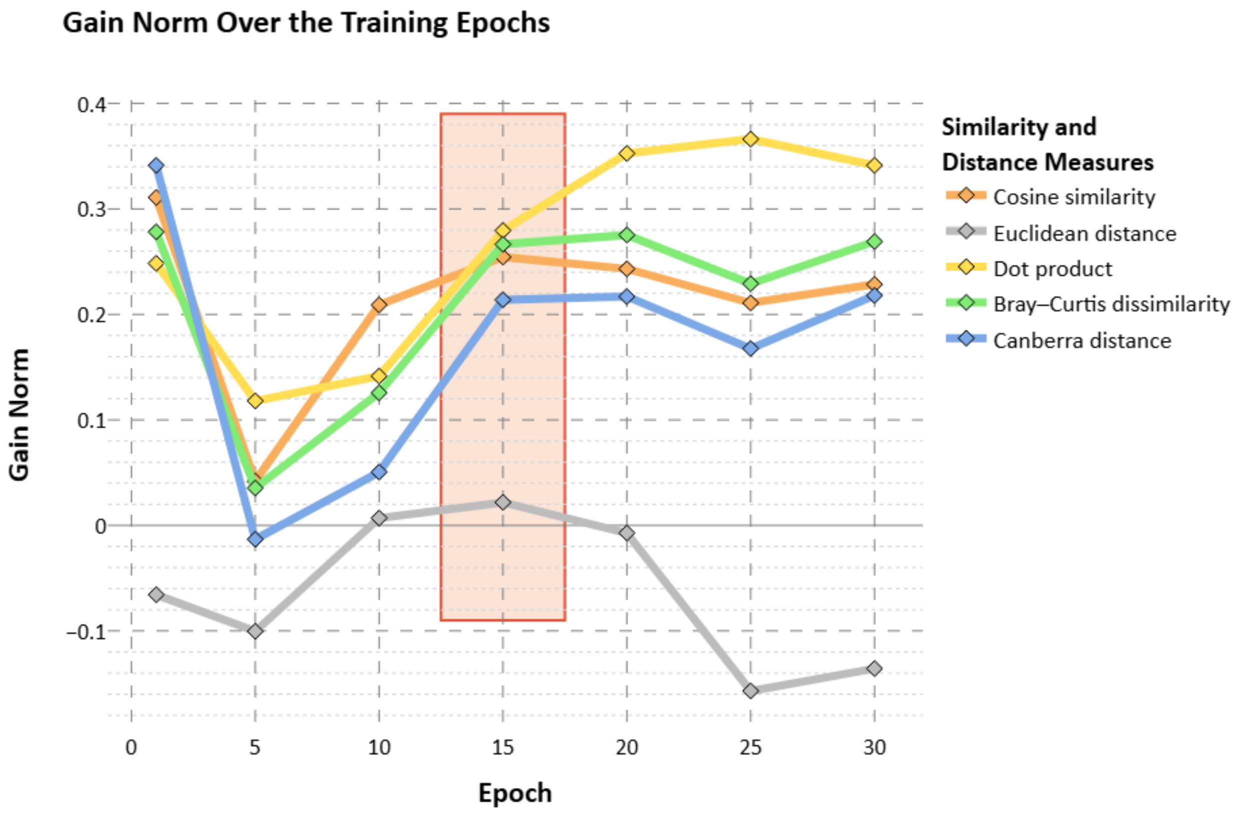

***Supplementary Figure 7.*** *Gain norm calculated with various similarity and distance measures throughout the training of the NSG model with reaction2all context-sampling. The epoch from which the final embedding was extracted is highlighted by a semi-transparent red rectangle.*

### 6.3. NTX-reaction2drug

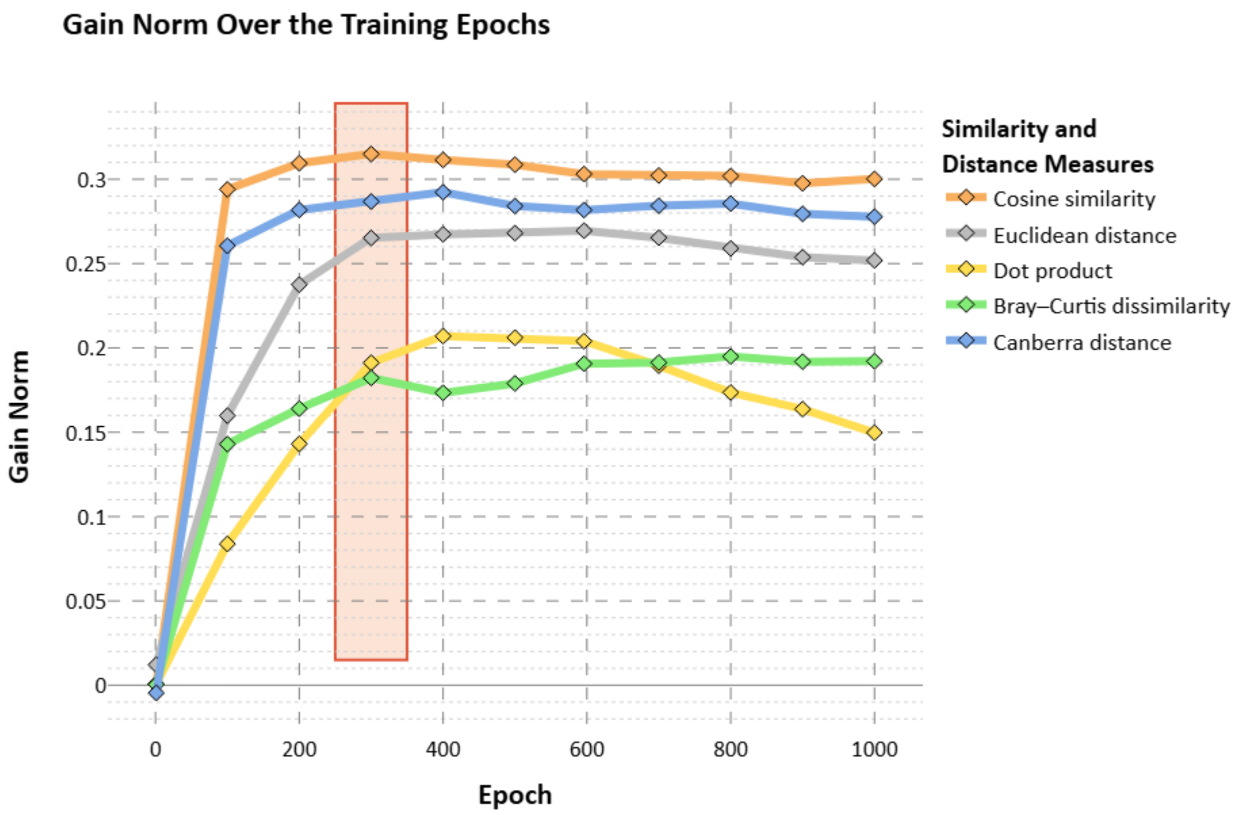

***Supplementary Figure 8.*** *Gain norm calculated with various similarity and distance measures throughout the training of the NTX model with reaction2drug context-sampling. The epoch from which the final embedding was extracted is highlighted by a semi-transparent red rectangle.*

### 6.4. NTX-reaction2reaction

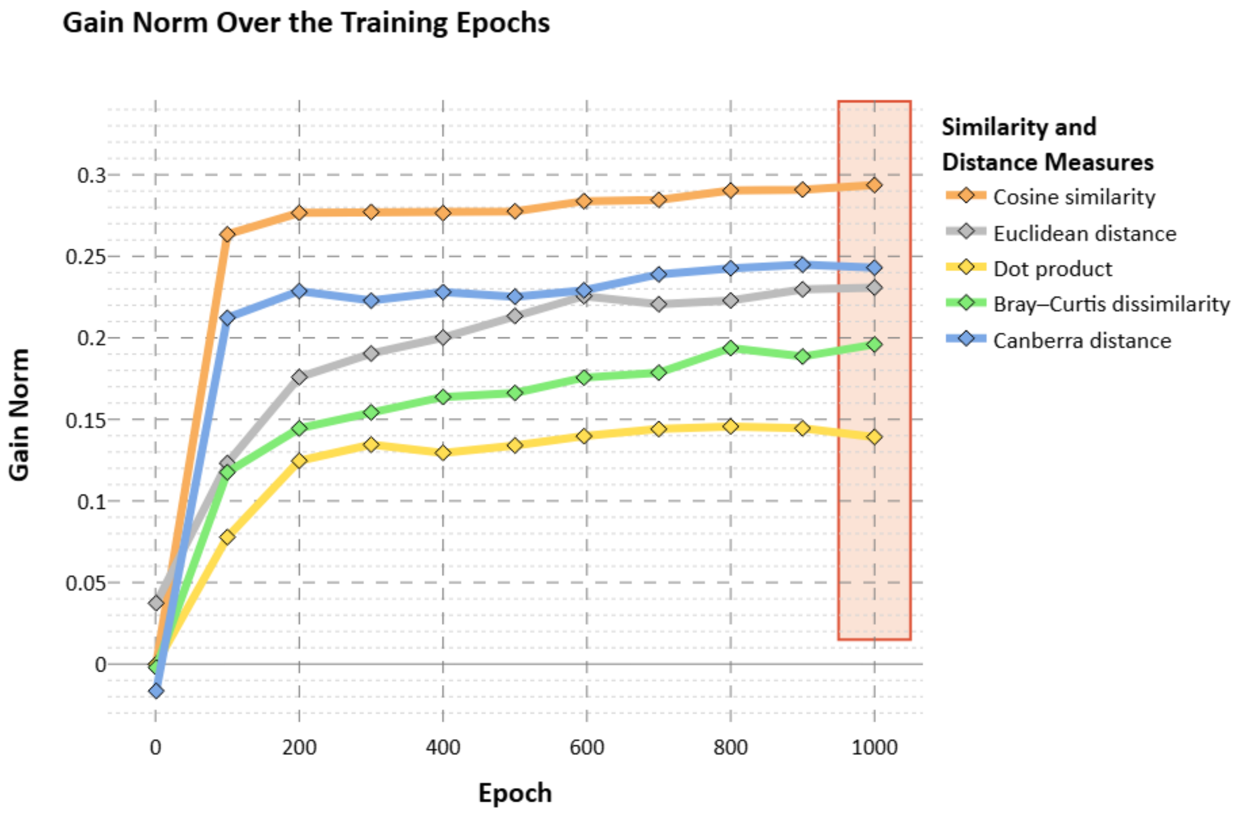

***Supplementary Figure 9.*** *Gain norm calculated with various similarity and distance measures throughout the training of the NTX model with reaction2reaction context-sampling. The epoch from which the final embedding was extracted is highlighted by a semi-transparent red rectangle.*

### 6.5. NTX-reaction2all

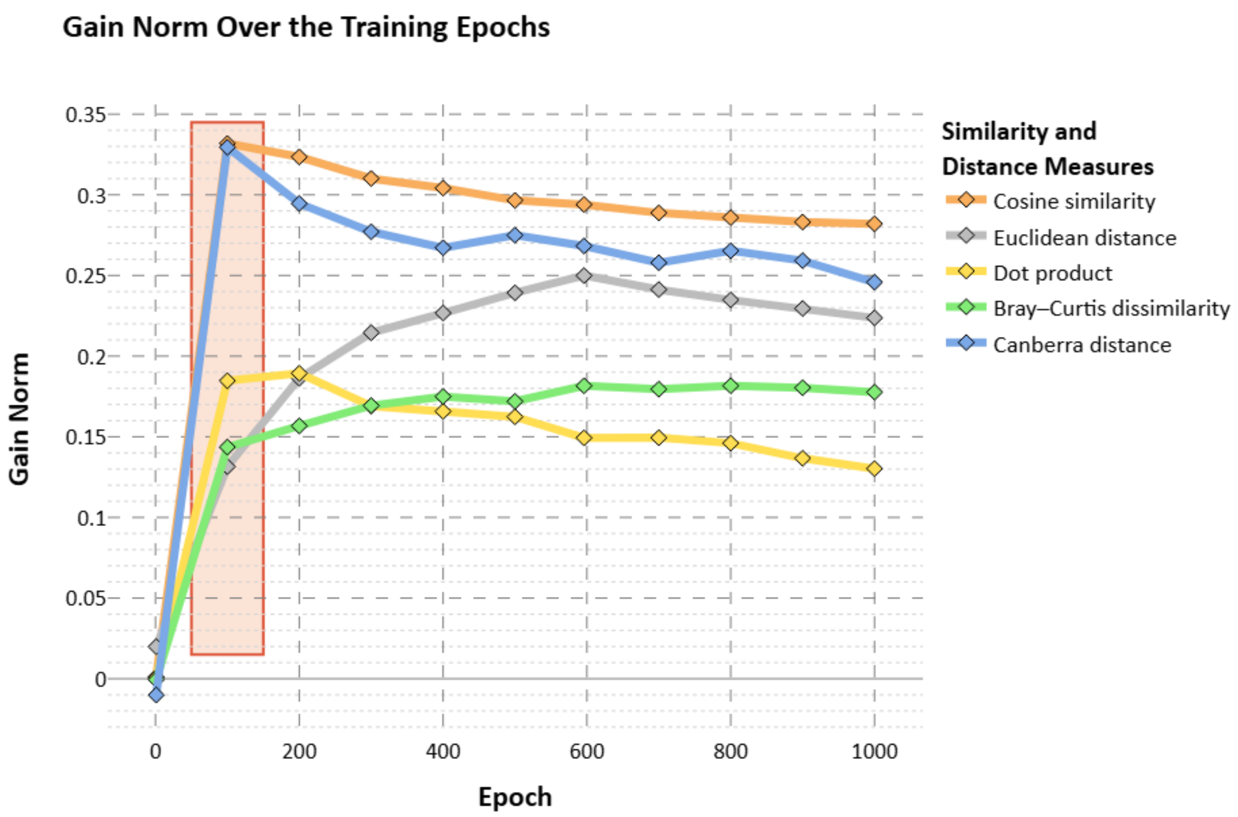

***Supplementary Figure 10.*** *Gain norm calculated with various similarity and distance measures throughout the training of the NTX model with reaction2all context-sampling. The epoch from which the final embedding was extracted is highlighted by a semi-transparent red rectangle.*

### 7. Classifier performance comparison

***Supplementary Table 10.*** *Performance of the ROR-based signal detection method and our classifier with the six different embeddings as reaction features, with the six different negative samplings, tested on benchmark datasets from three sources (here referred to as EU-ADR[18], OMOP[19] and FDA-rev[20]). Results in Table 1 of the manuscript are highlighted by a blue frame.*

| Negative sampling | | Prediction method | EU-ADR | | | OMOP | | | FDA-rev | | | Merged | | |
|---|---|---|---|---|---|---|---|---|---|---|---|---|---|---|
| Keeping | Sampling | | Acc | AUROC | AUPRC | Acc | AUROC | AUPRC | Acc | AUROC | AUPRC | Acc | AUROC | AUPRC |
| - | - | ROR | 0.8158 | 0.9094 | 0.9021 | 0.6867 | 0.7076 | 0.6695 | 0.6535 | **0.7592** | **0.7328** | 0.7071 | 0.7548 | 0.7357 |
| Drugs | Reactions with frequency | NSG-reaction2drug | 0.8110 | 0.8915 | 0.8738 | 0.7512 | 0.8159 | 0.7616 | 0.6632 | 0.7303 | 0.6509 | 0.7438 | 0.8094 | 0.7501 |
| | | NSG-reaction2reaction | 0.8211 | 0.8875 | 0.8498 | 0.7449 | 0.8242 | 0.7831 | 0.6440 | 0.7011 | 0.6003 | 0.7372 | 0.8081 | 0.7414 |
| | | NSG-reaction2all | 0.8147 | 0.8973 | 0.8816 | 0.7557 | 0.8478 | 0.7995 | 0.6648 | 0.7381 | 0.6386 | 0.7472 | 0.8324 | 0.7717 |
| | | NTX-reaction2drug | 0.7495 | 0.8207 | 0.8122 | 0.6288 | 0.6487 | 0.5685 | 0.6336 | 0.6962 | 0.5928 | 0.6525 | 0.6944 | 0.6180 |
| | | NTX-reaction2reaction | 0.7807 | 0.8552 | 0.8615 | 0.6418 | 0.6729 | 0.6083 | 0.6560 | 0.7001 | 0.6082 | 0.6708 | 0.7134 | 0.6507 |
| | | NTX-reaction2all | 0.7422 | 0.8182 | 0.8225 | 0.6244 | 0.6569 | 0.5361 | 0.6096 | 0.6654 | 0.5837 | 0.6431 | 0.6898 | 0.6030 |
| Reactions | Drugs with frequency | NSG-reaction2drug | 0.8339 | **0.9310** | **0.9187** | 0.8019 | 0.8717 | 0.8128 | 0.6784 | 0.7482 | 0.6852 | 0.7813 | 0.8551 | 0.7999 |
| | | NSG-reaction2reaction | **0.8541** | 0.9187 | 0.9127 | 0.8388 | 0.9167 | 0.8838 | 0.6688 | 0.7329 | 0.6662 | **0.8056** | 0.8775 | 0.8322 |
| | | NSG-reaction2all | 0.8358 | 0.9213 | 0.9133 | 0.8291 | 0.9098 | 0.8660 | **0.6808** | 0.7544 | 0.7000 | 0.7986 | **0.8801** | **0.8373** |
| | | NTX-reaction2drug | 0.8046 | 0.8915 | 0.8684 | 0.7036 | 0.7587 | 0.6665 | 0.6736 | 0.7376 | 0.6691 | 0.7153 | 0.7791 | 0.7013 |
| | | NTX-reaction2reaction | 0.8413 | 0.8972 | 0.9030 | 0.7222 | 0.7957 | 0.7303 | 0.6656 | 0.7175 | 0.6596 | 0.7312 | 0.7968 | 0.7424 |
| | | NTX-reaction2all | 0.7670 | 0.8486 | 0.8182 | 0.7291 | 0.7990 | 0.7036 | 0.6608 | 0.7041 | 0.6411 | 0.7222 | 0.7863 | 0.7010 |
| None | Both with frequency | NSG-reaction2drug | 0.7972 | 0.8987 | 0.8959 | 0.7657 | 0.8319 | 0.7787 | 0.6632 | 0.7240 | 0.6398 | 0.7492 | 0.8182 | 0.7580 |
| | | NSG-reaction2reaction | 0.8119 | 0.8971 | 0.8770 | 0.7898 | 0.8761 | 0.8433 | 0.6280 | 0.6915 | 0.6026 | 0.7591 | 0.8357 | 0.7787 |
| | | NSG-reaction2all | 0.8183 | 0.9102 | 0.9043 | 0.7884 | 0.8860 | 0.8447 | 0.6640 | 0.7273 | 0.6443 | 0.7674 | 0.8531 | 0.8026 |
| | | NTX-reaction2drug | 0.7651 | 0.8423 | 0.8339 | 0.6878 | 0.7453 | 0.6701 | 0.6344 | 0.6853 | 0.5800 | 0.6913 | 0.7505 | 0.6683 |
| | | NTX-reaction2reaction | 0.7560 | 0.8548 | 0.8686 | 0.6895 | 0.7466 | 0.7018 | 0.6400 | 0.6860 | 0.5995 | 0.6905 | 0.7484 | 0.6924 |
| | | NTX-reaction2all | 0.7239 | 0.8175 | 0.8094 | 0.6762 | 0.7584 | 0.6711 | 0.5888 | 0.6524 | 0.5720 | 0.6669 | 0.7421 | 0.6552 |
| Drugs | Reactions uniformly | NSG-reaction2drug | 0.5945 | 0.7501 | 0.7031 | 0.5452 | 0.6513 | 0.6020 | 0.6032 | 0.6560 | 0.5745 | 0.5637 | 0.6690 | 0.5973 |
| | | NSG-reaction2reaction | 0.5835 | 0.7598 | 0.7072 | 0.5313 | 0.7054 | 0.6823 | 0.5792 | 0.6160 | 0.5300 | 0.5493 | 0.6947 | 0.6223 |
| | | NSG-reaction2all | 0.5807 | 0.7583 | 0.7373 | 0.5427 | 0.6905 | 0.6438 | 0.5912 | 0.6439 | 0.5388 | 0.5576 | 0.6903 | 0.6129 |
| | | NTX-reaction2drug | 0.5312 | 0.6748 | 0.6725 | 0.4983 | 0.5711 | 0.4856 | 0.5688 | 0.6069 | 0.5128 | 0.5173 | 0.5975 | 0.5193 |
| | | NTX-reaction2reaction | 0.5587 | 0.7105 | 0.7072 | 0.5338 | 0.5898 | 0.5351 | 0.5544 | 0.6096 | 0.5359 | 0.5413 | 0.6154 | 0.5498 |
| | | NTX-reaction2all | 0.5413 | 0.6641 | 0.5951 | 0.5075 | 0.5882 | 0.4966 | 0.5488 | 0.5794 | 0.5096 | 0.5209 | 0.5991 | 0.5122 |
| Reactions | Drugs uniformly | NSG-reaction2drug | 0.7578 | 0.8869 | 0.8703 | 0.8144 | 0.8945 | 0.8521 | 0.6488 | 0.6967 | 0.6255 | 0.7684 | 0.8515 | 0.7984 |
| | | NSG-reaction2reaction | 0.7495 | 0.8652 | 0.8447 | **0.8460** | **0.9215** | **0.8902** | 0.6488 | 0.6924 | 0.6340 | 0.7857 | 0.8636 | 0.8140 |
| | | NSG-reaction2all | 0.7422 | 0.8759 | 0.8717 | 0.8400 | 0.9142 | 0.8746 | 0.6616 | 0.7026 | 0.6559 | 0.7839 | 0.8645 | 0.8225 |
| | | NTX-reaction2drug | 0.7394 | 0.8419 | 0.8213 | 0.7463 | 0.8179 | 0.7506 | 0.6320 | 0.6618 | 0.5908 | 0.7194 | 0.7873 | 0.7185 |
| | | NTX-reaction2reaction | 0.7312 | 0.8333 | 0.8290 | 0.7615 | 0.8427 | 0.7815 | 0.6344 | 0.6745 | 0.6018 | 0.7282 | 0.7985 | 0.7276 |
| | | NTX-reaction2all | 0.6670 | 0.7734 | 0.7529 | 0.7706 | 0.8495 | 0.7909 | 0.5960 | 0.6274 | 0.5721 | 0.7138 | 0.7857 | 0.7126 |
| None | Both uniformly | NSG-reaction2drug | 0.6037 | 0.7897 | 0.7673 | 0.6227 | 0.7889 | 0.7421 | 0.5912 | 0.6385 | 0.5449 | 0.6098 | 0.7528 | 0.6815 |
| | | NSG-reaction2reaction | 0.6055 | 0.7800 | 0.7203 | 0.6377 | 0.8144 | 0.7895 | 0.5608 | 0.6049 | 0.5225 | 0.6127 | 0.7610 | 0.6860 |
| | | NSG-reaction2all | 0.5972 | 0.7917 | 0.7673 | 0.6488 | 0.8122 | 0.7668 | 0.5936 | 0.6377 | 0.5386 | 0.6255 | 0.7682 | 0.6908 |
| | | NTX-reaction2drug | 0.5817 | 0.7034 | 0.6830 | 0.5584 | 0.5953 | 0.5010 | 0.5823 | 0.7310 | 0.6761 | 0.5745 | 0.6923 | 0.6049 |
| | | NTX-reaction2reaction | 0.6037 | 0.7338 | 0.7059 | 0.6285 | 0.7574 | 0.7090 | 0.5736 | 0.6241 | 0.5251 | 0.6100 | 0.7192 | 0.6310 |
| | | NTX-reaction2all | 0.5523 | 0.6761 | 0.6074 | 0.6019 | 0.7506 | 0.7091 | 0.5336 | 0.5761 | 0.4946 | 0.5759 | 0.6952 | 0.5979 |

*ROR: reporting odds ratio; Acc: accuracy; AUROC: area under the receiving operating characteristic curve; AUPRC: area under the precision-recall curve*

### 8. The effects of negative sampling

(**Supplementary Discussion 1**)

The results shown in Table 1 of the manuscript (highlighted by a blue frame in Supplementary Table 10) are from the negative sampling approach where we keep the reactions and sample new drugs by their frequency distribution as seen in the validated data. We observed that the main contributor to the varying performance of the classifier was whether we sampled uniformly or by frequency, as that significantly influenced the distribution differences of the constituents. Sampling uniformly made it too easy for the classifier to separate the uniform negatives from the heavily skewed positives, which resulted in high performance metrics during training but low values during testing. Matching the frequency of the constituents created a more balanced dataset and an overall harder training task, leading to better generalization capabilities.

For this reason, we hypothesized that the best performing sampling method will be when none of the constituents are kept and instead both are resampled with their corresponding frequency distribution (last group of Supplementary Table 10) but that turned out to be false. With keeping the reactions and sampling the drugs by their frequency, the distributions of reactions on the positive and negative sides are the exact same, as all reactions have an equal amount of positive and negative drug counterparts. This forces the model even stronger not to ignore the drugs, and instead consider how the reaction embedding vectors and the drug feature vectors interact. With the other way around, when the drugs are kept and the reactions are sampled by their frequency, we observed worse performances compared to the other two frequency-based approaches, suggesting that the reaction embedding vectors are more information rich than the drug feature vectors and could overshadow their contribution when allowed to.

We might also mention an alternative approach of constructing negative data, namely to label all drug–event pairs from the report corpus as negative if they are not present in the validated datasets. This option would result in a training objective where the model is expected to differentiate pairs that were recorded in the text-mined sources of the validated datasets from pairs that were reported but not recorded. We considered this approach to be theoretically unfound, as there could be several aspects, other than causality, which influenced whether a pair was included in the validated data or only in the reports, resulting in a model that might fit to these distinguishing patterns instead. For example, the validated data might overrepresent pairs with trivial associations, or drugs that simply had more time on the market, or adverse events that are of great health concern and enjoy more attention from authorities and researchers. These potential issues are mostly addressed by our frequency-based negative sampling approaches.

### 9. Assessment of drug–event misclassifications by the best performing model

***Supplementary Table 11.*** *Assessment of the greatest misclassifications (predicted confidence is either* $> 0.75$ *or* $< 0.25$*) on the benchmark sets from various sources (here referred to as EU-ADR[18], OMOP[19] and FDA-rev[20]) by our best performing classifier model using the NSG-reaction2all embedding (see Supplementary Discussion 1). Comparison is made against information in the available literature to determine whether the truth of the benchmark sets or the prediction of the classifier is more correct about the actual causality between the given drug and reaction.*

| Drug | Reaction | Source | Truth | Model | Causality | Assessment |
|---|---|---|---|---|---|---|
| Methylphenidate | Nightmare | FDA-rev | 0 | 0.8859 | **Likely** | Sleep disturbance has a significant relationship with *Methylphenidate*, and therefore *Nightmare* could be a possible reason.[21] |
| Sodium Phosphate, Monobasic | Acute Myocardial Infarction | OMOP | 0 | 0.8570 | **Likely** | Although the overdose of *Sodium Phosphate* induces myocardial necrosis in rats, conclusive evidence in human studies could not be found for obvious reasons.[22] |
| Doxazosin | Rhabdomyolysis | EU-ADR | 0 | 0.8407 | **No evidence** | There is no evidence on the connection between *Doxazosin* and *Rhabdomyolysis*. |
| Sodium Phosphate, Monobasic | Acute Kidney Injury | OMOP | 0 | 0.8294 | **Controversial evidence** | *Sodium Phosphate* may contribute to *Acute Kidney Injury* if other drugs are present. As an independent reaction, it is debated in the literature.[23,24] |
| Rilpivirine | Drug Reaction with Eosinophilia and Systemic Symptoms | FDA-rev | 0 | 0.8243 | **No evidence** | *Eosinphilia and Systemic Symptoms* are well-known signs of allergy, and drug allergy is quite common.[25] |
| Levodopa | Acute Kidney Injury | EU-ADR | 0 | 0.8008 | **Unlikely** | There is clear evidence that *Levodopa* administration does not induce *Acute Kidney Injury*. *Levodopa* is a precursor of dopamine which has renal effects.[26] |
| Darunavir | Acute Kidney Injury | OMOP | 0 | 0.7928 | **Unlikely** | There is no comprehensive study concerning this topic, only case studies with COVID19 involved, which might cause *Acute Kidney Injury* by itself. |
| Sotalol | Agranulocytosis | EU-ADR | 0 | 0.7861 | **Unlikely** | Though other antiarrhythmic agents co-administered with *Sotalol*, like amiodarone, might cause aplastic anemia, there is no evidence that *Sotalol* causes *Agranulocytosis*.[27] |
| Pantoprazole | Dyspnoea | FDA-rev | 0 | 0.7756 | **Unlikely** | *Dyspnoea* is a common symptom of cardiovascular diseases. In cardiac care, antithrombotics are frequently used, which may contribute to development of gastric ulcers. Therefore, in general practice, *Pantoprazole* is usually co-administered with antithrombotic drugs as a preventive measure against ulcers.[2] Thus, *Dyspnoea* might correlate with *Pantoprazole*, while their direct relation seems to be untrue. |

| | | | | | | |
|---|---|---|---|---|---|---|
| Ferrous Gluconate | Acute Kidney Injury | OMOP | 0 | 0.7742 | **No evidence** | Even though chronic renal diseases have connections with iron, via eritropoetin deficiency and microcytic anemia, there is no evidence suggesting the causality between *Ferrous Gluconate* and *Acute Kidney Injury*.[28] |
| Desipramine | Acute Myocardial Infarction | OMOP | 1 | 0.2450 | **Controversial evidence** | Even though rat studies state that *Desipramine* reduces infarction size through regulating norepinephrine release, we did not find any studies stating the same in humans.[29] |
| Ponatinib | Ocular Toxicity | FDA-rev | 1 | 0.2412 | **Clear evidence** | Several studies suggest ocular adverse drug reactions, advising regular ophthalmological checkups.[30] |
| Levosalbutamol | Gastrooesophageal Reflux Disease | FDA-rev | 1 | 0.2276 | **Clear evidence** | *Levosalbutamol* is an enantiomer of salbutamol (or alternatively albuterol), which inhibits esophageal motion. There is no study that suggest the other enantiomer would cause the side effect and the cited study shows that the intravenous form of salbutamol causes *Gastrooesophageal Reflux Disease*.[31] |
| Rosiglitazone | Acute Myocardial Infarction | EU-ADR | 1 | 0.2170 | **Clear evidence** | Several studies state that *Rosiglitazone* does contribute to *Acute Myocardial Infarction*.[32] |
| Amoxapine | Acute Myocardial Infarction | OMOP | 1 | 0.2032 | **Unlikely** | The cited study shows that there is a slight decrease in the risk of *Acute Myocardial Infarction* as opposed to not taking a selective serotonin reuptake inhibitor, like *Amoxapine*.[33] |
| Lisdexamfetamine | Peripheral Vascular Disorder | FDA-rev | 1 | 0.1578 | **Contra-indication** | *Peripheral Vascular Disorder* is listed as a **contra-indication** for amphetamine substances, meaning *Lisdexamfetamine* does not necessarily cause *Peripheral Vascular Disorder* but might worsen the condition of the patient having it.[34] |
| Sulfamethoxazole | Electrocardiogram QT Prolonged | FDA-rev | 1 | 0.1459 | **Clear evidence** | Several studies state that *Sulfamethoxazole* does contribute to *Electrocardiogram QT Prolonged*.[35] |
| Dextroamphetamine | Peripheral Vascular Disorder | FDA-rev | 1 | 0.0673 | **Contra-indication** | See *Lisdexamfetamine*. |
| Metamfetamine | Peripheral Vascular Disorder | FDA-rev | 1 | 0.0360 | **Contra-indication** | See *Lisdexamfetamine*. |

### 10. References

1. Bill, R. W. *et al.* Evaluating semantic relatedness and similarity measures with Standardized MedDRA Queries. in *AMIA Annual Symposium Proceedings* vol. 2012 43–50 (American Medical Informatics Association, 2012).
2. Byrne, R. A. *et al.* 2023 ESC Guidelines for the management of acute coronary syndromes: Developed by the task force on the management of acute coronary syndromes of the European Society of Cardiology (ESC). *Eur. Heart J.* **44**, 3720–3826 (2023).
3. Vrints, C. *et al.* 2024 ESC Guidelines for the management of chronic coronary syndromes: Developed by the task force for the management of chronic coronary syndromes of the European Society of Cardiology (ESC) Endorsed by the European Association for Cardio-Thoracic Surgery. *Eur. Heart J.* **45**, 3415–3537 (2024).
4. Thygesen, K. *et al.* Fourth universal definition of myocardial infarction (2018). *Eur. Heart J.* **40**, 237–269 (2019).
5. Stevens, P. E. *et al.* KDIGO 2024 Clinical Practice Guideline for the Evaluation and Management of Chronic Kidney Disease. *Kidney Int.* **105**, S117–S314 (2024).
6. European Association for the Study of the Liver (EASL). EASL Clinical Practice Guidelines on the prevention, diagnosis and treatment of gallstones. *J. Hepatol.* **65**, 146–181 (2016).
7. American Diabetes Association Professional Practice Committee. 6. Glycemic Goals and Hypoglycemia: Standards of Care in Diabetes—2025. *Diabetes Care* **48**, S128–S145 (2024).
8. McMurray, J. *et al.* Kidney disease: Improving global outcomes (KDIGO) anemia work group. KDIGO clinical practice guideline for anemia in chronic kidney disease. *Kidney Int. Suppl.* 279–335 (2012).
9. American Diabetes Association Professional Practice Committee. 12. Retinopathy, Neuropathy, and Foot Care: Standards of Care in Diabetes—2025. *Diabetes Care* **48**, S252–S265 (2024).
10. American Diabetes Association Professional Practice Committee. 2. Diagnosis and Classification of Diabetes: Standards of Care in Diabetes—2025. *Diabetes Care* **48**, S27–S49 (2024).
11. American Diabetes Association Professional Practice Committee. 13. Older Adults: Standards of Care in Diabetes—2025. *Diabetes Care* **48**, S266–S282 (2024).
12. Anderson, M. J. Distance-based tests for homogeneity of multivariate dispersions. *Biometrics* **62**, 245–253 (2006).
13. Gijbels, I. & Omelka, M. Testing for Homogeneity of Multivariate Dispersions Using Dissimilarity Measures. *Biometrics* **69**, 137–145 (2013).
14. Benjamini, Y. & Hochberg, Y. Controlling the False Discovery Rate: A Practical and Powerful Approach to Multiple Testing. *J. R. Stat. Soc. Ser. B Stat. Methodol.* **57**, 289–300 (1995).
15. Cailliez, F. The analytical solution of the additive constant problem. *Psychometrika* **48**, 305–308 (1983).
16. Legendre, P. & Legendre, L. Numerical Ecology. in *Encyclopedia of Ecology: Volume 1-4, Second Edition* vol. 3 487–493 (Elsevier, 2019).

17. Vardi, Y. & Zhang, C. H. The multivariate L1-median and associated data depth. in *Proceedings of the National Academy of Sciences of the United States of America* vol. 97 1423–1426 (2000).

18. Coloma, P. M. *et al.* A reference standard for evaluation of methods for drug safety signal detection using electronic healthcare record databases. *Drug Saf.* **36**, 13–23 (2013).

19. Ryan, P. B. *et al.* Defining a reference set to support methodological research in drug safety. *Drug Saf.* **36**, 33–47 (2013).

20. Harpaz, R. *et al.* A time-indexed reference standard of adverse drug reactions. *Sci. Data* **1**, 1–10 (2014).

21. Storebø, O. J. *et al.* Methylphenidate for children and adolescents with attention deficit hyperactivity disorder (ADHD). *Cochrane Database Syst. Rev.* **2023**, (2023).

22. Lehr, D. & Krukowski, M. About the Mechanism of Myocardial Necrosis Induced by Sodium Phosphate and Adrenal Corticoid Overdosage. *Ann. N. Y. Acad. Sci.* **105**, 137–182 (1963).

23. Layton, J. B. *et al.* Sodium phosphate does not increase risk for acute kidney injury after routine colonoscopy, compared with polyethylene glycol. *Clin. Gastroenterol. Hepatol.* **12**, (2014).

24. Markowitz, G. S. & Perazella, M. A. Acute phosphate nephropathy. *Kidney Int.* **76**, 1027–1034 (2009).

25. Tebas, P. *et al.* Lipid Levels and Changes in Body Fat Distribution in Treatment-Naive, HIV-1–Infected Adults Treated With Rilpivirine or Efavirenz for 96 Weeks in the ECHO and THRIVE Trials. *Clin. Infect. Dis.* **59**, 425–434 (2014).

26. Boelens Keun, J. T., Arnoldussen, I. A., Vriend, C. & Van De Rest, O. Dietary Approaches to Improve Efficacy and Control Side Effects of Levodopa Therapy in Parkinson's Disease: A Systematic Review. *Adv. Nutr.* **12**, 2265–2287 (2021).

27. Somberg, J. & Molnar, J. Sotalol versus Amiodarone in Treatment of Atrial Fibrillation. *J. Atr. Fibrillation* **8**, 1359 (2016).

28. Cancelo-Hidalgo, M. J. *et al.* Tolerability of different oral iron supplements: a systematic review. *Curr. Med. Res. Opin.* **29**, 291–303 (2013).

29. Borganelli, M. & Forman, M. B. Simulation of acute myocardial infarction by desipramine hydrochloride. *Am. Heart J.* **119**, 1413–1414 (1990).

30. Moon, J. A., Bowden, E. C. & Aung, M. H. Probable ponatinib-induced papilledema in a patient with acute lymphocytic leukemia. *Am. J. Ophthalmol. Case Reports* **32**, (2023).

31. Schindlbeck, N. E., Heinrich, C., Huber, R. M. & Müller Lissner, S. A. Effects of Albuterol (Salbutamol) on Esophageal Motility and Gastroesophageal Reflux in Healthy Volunteers. *JAMA* **260**, 3156–3158 (1988).

32. Kroker, A. J. & Bruning, J. B. Review of the Structural and Dynamic Mechanisms of PPARγ Partial Agonism. *PPAR Res.* **2015**, 816856 (2015).

33. Schlienger, R. G., Fischer, L. M., Jick, H. & Meier, C. R. Current use of selective serotonin reuptake inhibitors and risk of acute myocardial infarction. *Drug Saf.* **27**, 1157–1165 (2004).

34. Tan, G. M. *et al.* Peripheral vascular manifestation in patients receiving an amphetamine analog: A case series. *Vasc. Med. (United Kingdom)* **24**, 50–55 (2019).

35. Lopez, J. A. *et al.* QT prolongation and torsades de pointes after administration of trimethoprim-sulfamethoxazole. *Am. J. Cardiol.* **59**, 376–377 (1987).
